# HapBridge: A Methylation-Guided Approach for Correcting Switch Errors and Bridging Phased Blocks in Long-Read Phasing

**DOI:** 10.1101/2025.11.07.687303

**Authors:** Hansen Chen, Fan Nie, Feng Luo, Jianxin Wang

**Affiliations:** School of Computer Science and Engineering, Central South University, Changsha, 410083, China; Xiangjiang Laboratory, Changsha, 410205, China; Hunan Provincial Key Lab on Bioinformatics, Central South University, Changsha, 410083, China; School of Mathematics and Computational Science, Xiangtan University, Xiangtan, 41105, China; National Center for Applied Mathematics in Hunan and Key Laboratory of Intelligent Computing and Information Processing of Ministry of Education, Xiangtan University, Xiangtan, Hunan 41105, China; School of Computing, Clemson University, Clemson, SC, 29634-0974, USA

**Author notes:** These authors contributed equally: Hansen Chen, Fan Nie. To whom correspondence should be addressed: Jianxin Wang.

**Keywords:** DNA sequencing, haplotype phasing, methylation

## Abstract

Long-read sequencing has substantially advanced haplotype phasing yet continues to face challenges in low-heterozygosity regions and switch errors caused by read noise. DNA methylation exhibits haplotype-specific patterns, providing complementary linkage information, but existing phasing algorithms have not fully leveraged these signals owing to their variability. Here, we present **HapBridge**, a methylation-guided phasing framework that enhances single-nucleotide variant (SNV)-based phasing by **detecting and correcting switch errors** and **bridging adjacent phased blocks**. Evaluations on Oxford Nanopore R9/R10 and PacBio HiFi datasets show that HapBridge reduces switch errors by **3.07–18.72%** and improves N50 length by **5.84–68.61%** compared with MethPhaser, achieving higher phasing accuracy and contiguity. These results demonstrate that integrating methylation with sequence variation can provide a robust and intrinsic linkage signal that effectively improves haplotype continuity in long-read sequencing. **HapBridge** is publicly available at GitHub https://github.com/Humonex/HapBridge.

## 1 Introduction

In diploid organisms, which inherit one homologous chromosome set from each parent, the assignment of alleles at heterozygous sites to their corresponding haplotypes is referred to as phasing. Accurate reconstruction of maternal and paternal haplotypes is essential for elucidating inheritance patterns, regulatory mechanisms, and the effects of genetic variation on various traits and diseases [9,37,29,30].

Current phasing algorithms typically use the linkage information among shared variants within the same sequencing read to assign alleles. However, due to inherent read-length constraints, these methods usually produce localized phasing (discrete phased blocks), leaving unphased gaps between blocks where linkage information is absent.

Recent advances in sequencing technologies, particularly long-read platforms [11], have enabled the identification of genetic variations, such as single-nucleotide polymorphisms (SNPs) [36,13], insertions and deletions (InDels) [26], and structural variants (SVs) [34,6]. Variant calling tools—such as DeepVariant [24], GATK [25], and Clair3 [39]—have been developed for single nucleotide variant (SNV) detection and achieve consistently high accuracy across diverse sequencing platforms. Based on these developments, SNV-based phasing tools such as WhatsHap [19], HapCUT2 [5], LongPhase [18], and MARGIN [4] utilize long-read sequencing data to phase haplotypes. WhatsHap, a read-based phasing tool, solves the weighted minimum error correction (wMEC) problem to infer haplotypes from reads spanning multiple heterozygous variants. HapCUT2 uses a likelihood-based error-correction framework for constructing phased blocks. MARGIN integrates a hidden Markov model (HMM) to perform haplotagging and variant phasing on long reads. LongPhase simultaneously co-phasing SNPs, small indels, and structural variants in the human genome. Although these methods perform well in regions with abundant SNVs, they often fail to phase across regions of low heterozygosity or sparsely distributed variants. To address the limitations of conventional phasing methods in regions with low heterozygosity or sparsely distributed variants, complementary data sources such as Hi-C [17,35,16], parental genotypes [12], and RNA sequencing reads [2,12] have been increasingly utilized to enhance phasing accuracy and continuity. While those methods can enhance phasing performance—sometimes achieving chromosome-scale haplotypes—their reliance on additional sequencing data poses practical limitations. The availability and cost of such data significantly constrain their broad application.

Another promising direction for overcoming these limitations involves utilizing DNA methylation signals as an additional layer of information [20,33,31,21,38,10]. With the advancement of nanopore sequencing technologies, several computational frameworks, including Nanopolish [33], Remora [22], and Dorado [23], have been developed to detect 5mC at CpG sites, substantially improving both the sensitivity and accuracy of methylation calling. Recent progress in long-read sequencing and methylation detection has revealed that allele-specific methylation (ASM) strongly correlates with sequence haplotypes, offering an additional dimension for phasing beyond sequence variants alone [28,3,7,1]. Existing approaches, such as MethHaplo [40] and MethPhaser [8], have explored this potential, yet they remain limited. MethHaplo relies on bisulfite and Hi-C data, which limits its applicability, while MethPhaser, although fully utilizing long-read methylation signals, remains highly sensitive to the initial phasing accuracy and fails to detect or correct switch errors. Consequently, the integration of methylation information into phasing pipelines is still incomplete and suboptimal.

Here, we present HapBridge, a methylation-guided phasing framework that complements SNV-based phasers to achieve both switch error correction and block bridging in long-read datasets. HapBridge identifies haplotype-specific methylation loci and leverages their inter-site consistency to detect and correct switch errors within phased blocks, followed by an iterative bridging process that extends phasing continuity across previously unlinked regions. Unlike previous methods, HapBridge requires no additional sequencing data and can be seamlessly integrated into existing pipelines. Evaluations on Oxford Nanopore (ONT R9/R10) and PacBio HiFi datasets demonstrate that HapBridge consistently improves phasing accuracy and contiguity over state-of-the-art tools, confirming the value of methylation as an intrinsic, generalizable linkage signal for long-read haplotyping.

## 2 Results

### 2.1 Overview of the HapBridge

HapBridge processes the corresponding phased VCF and haplotagged BAM files generated by SNV-based phasers (e.g., HapCUT, WhatsHap; see **Methods**). It outputs refined phasing results, as shown in Figure 1a.

**Fig. 1.**
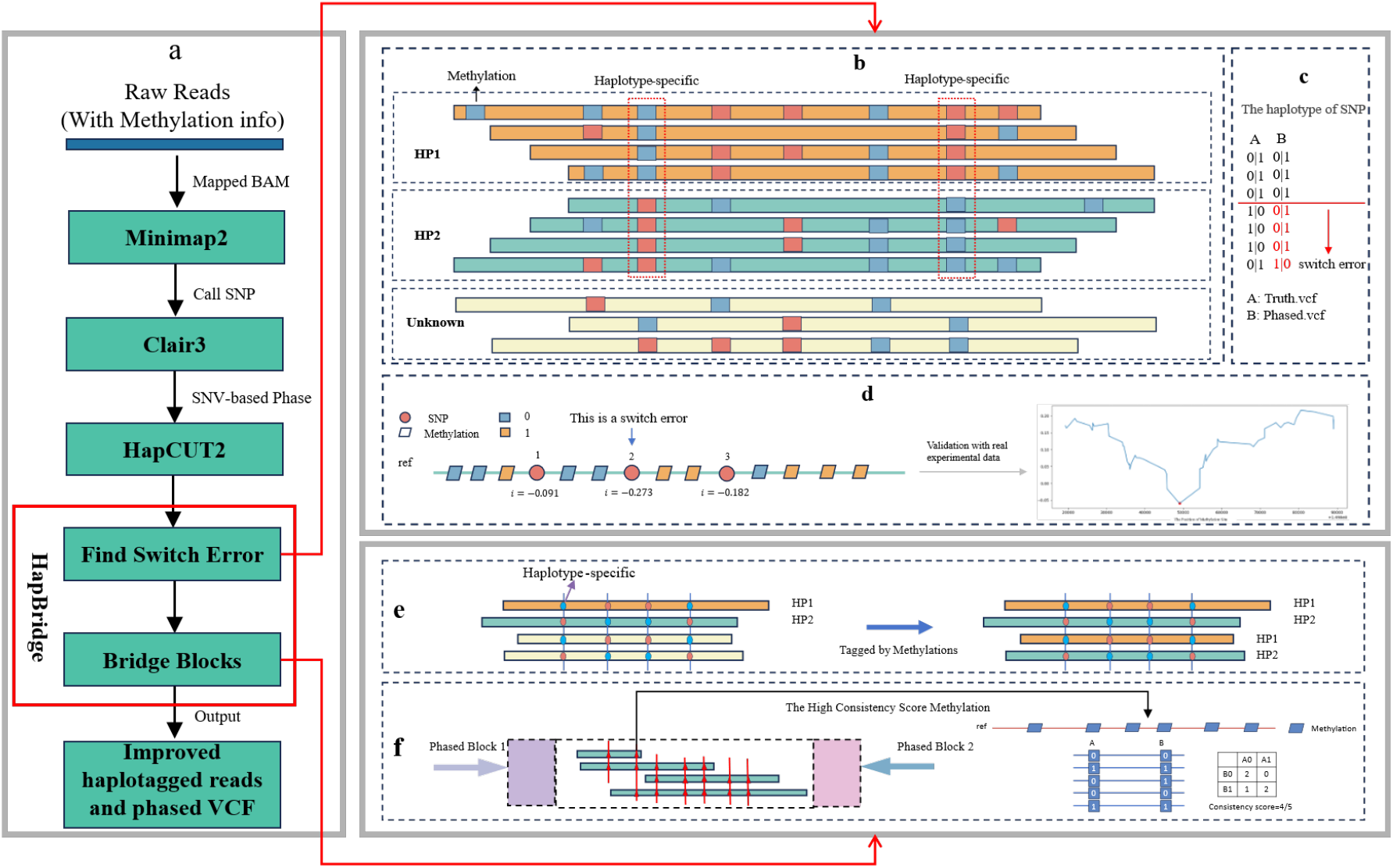
Overview of the HapBridge framework. HapBridge integrates methylation signals with SNV-based phasing to correct switch errors and bridge adjacent phase blocks. **a**, Workflow illustrating input BAM/VCF files, methylation extraction, switch-error correction, and iterative block bridging. **b–d**, Identification of haplotype-specific CpG sites and correction of local switch errors using methylation-consistency scores. **e–f**, Iterative expansion of phase blocks by retagging unphased reads and selecting high-confidence methylation anchors until convergence. HapBridge outputs an enhanced phased VCF and haplotagged BAM.

HapBridge consists of two main modules: *switch-error-correction* and *block-bridging*. The *switch-error-correction* module (Figure 1b-d) identifies haplotype-specific methylation sites and leverages their inter-site consistency to pinpoint and correct switch errors. The *block-bridging* module (Figure 1e-f) iteratively infers haplotype tags for unphased reads using haplotype-specific methylation sites and subsequently uses these reads to identify new haplotype-specific methylation sites. The iteration continues until no additional reads can be phased or no new methylation sites can be identified—or until it successfully bridges the two adjacent phased blocks. Throughout the iterations, methylation sites with high consistency scores are selected as anchors to connect adjacent phased blocks. A detailed description of HapBridge is provided in the **Methods** section.

### 2.2 Phasing performance evaluation

#### Switch error and flip Error performance evaluation

To comprehensively evaluate the phasing accuracy of HapBridge, we compared its performance against HapCUT2 and MethPhaser using two standard metrics: switch error and flip error [19](For clarity, HapCUT2-HapBridge and HapCUT2-MethPhaser denote that HapCUT2 was used as the underlying SNP phasing algorithm for HapBridge and Methphaser, respectively). These metrics quantify phasing accuracy across long-read data from the HG002 genome under different sequencing technologies (Oxford Nanopore R9/R10 and PacBio HiFi) and sequencing depths (80×, 60×, and 30×). Results were computed using the WhatsHap *compare* module (see Supplementary File 2 Section 4) and evaluated against the Genome in a Bottle (GIAB) SNP benchmark (v4.2.1).

As shown in Table 1, for ONT R9 data at high coverage (80×), HapCUT2-HapBridge achieves the lowest switch error (645), outperforming both HapCUT2 and HapCUT2-MethPhaser. At coverage (60×), HapCUT2-HapBridge maintains superior performance, with approximately 70 fewer switch errors compared to the other methods. Although at low coverage (30×), HapCUT2 exhibits a slight advantage over HapCUT2-HapBridge (36 fewer errors), but HapCUT2-HapBridge still achieves comparable accuracy. HapCUT2-MethPhaser consistently exhibits the highest switch-error rate.

**Table 1.**
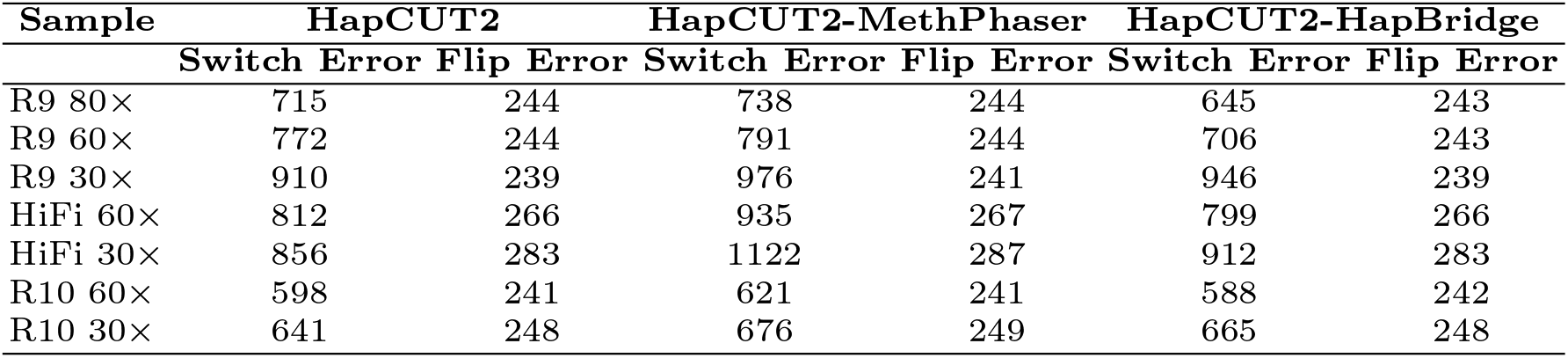
Comparison of Switch Error and Flip Error for HapCUT2, HapCUT2-HapBridge and HapCUT2-MethPhaser on the HG002 sample under different sequencing platforms (ONT R9/R10, PacBio HiFi) and read coverages (80×, 60×, 30×)

| Sample | HapCUT2 |  | HapCUT2-MethPhaser |  | HapCUT2-HapBridge |  |
| --- | --- | --- | --- | --- | --- | --- |
|  | Switch Error | Flip Error | Switch Error | Flip Error | Switch Error | Flip Error |
| R9 80× | 715 | 244 | 738 | 244 | 645 | 243 |
| R9 60× | 772 | 244 | 791 | 244 | 706 | 243 |
| R9 30× | 910 | 239 | 976 | 241 | 946 | 239 |
| HiFi 60× | 812 | 266 | 935 | 267 | 799 | 266 |
| HiFi 30× | 856 | 283 | 1122 | 287 | 912 | 283 |
| R10 60× | 598 | 241 | 621 | 241 | 588 | 242 |
| R10 30× | 641 | 248 | 676 | 249 | 665 | 248 |

**Table 2.** Independent contributions of the *switch-error-correction* and *block-bridging* modules across different sequencing platforms and coverages.

| Sample | HapCUT2 |  | switch-error-correction |  | block-bridging |  | HapBridge |  |
| --- | --- | --- | --- | --- | --- | --- | --- | --- |
|  | Switch Error | N50 | Switch Error | N50 | Switch Error | N50 | Switch Error | N50 |
| R9 80× | 715 | 2263840 | 642 | 2263840 | 718 | 4850574 | 645 | 4850574 |
| R9 60× | 772 | 2035463 | 655 | 2035463 | 823 | 2939026 | 706 | 2939026 |
| R10 60× | 598 | 2393002 | 576 | 2393002 | 610 | 3942177 | 588 | 3942177 |
| R10 30× | 641 | 1930657 | 615 | 1930657 | 691 | 2609119 | 665 | 2609119 |
| HiFi 30× | 856 | 414153 | 855 | 414153 | 913 | 507697 | 912 | 507697 |

For ONT R10 data, a similar trend is observed, with HapCUT2-HapBridge generally outperforming the other methods in reducing switch errors. For PacBio HiFi data, HapCUT2-HapBridge shows superior performance at 60× coverage, achieving 13 fewer switch errors than HapCUT2. At 30× coverage, HapCUT2 slightly outperforms HapCUT2-HapBridge; however, both methods substantially outperform HapCUT2-MethPhaser, highlighting the robustness of HapCUT2-HapBridge even under reduced coverage. Detailed results for each chromosome across all datasets are provided in the Supplementary File 2 Tables S1-S8.

Regarding flip errors, the results are nearly identical across all three methods, regardless of sequencing platform or coverage level, indicating that all tools offer similar precision for this metric. Taken together, these results demonstrate that HapCUT2-HapBridge remains highly competitive across technologies and sequencing depths, consistently delivering the lowest switch-error rates at medium to high coverage. Supplementary File 2 Tables S9-S17 show the switch error and flip error of WhatsHap, WhatsHap-MethPhaser, and WhatsHap-HapBridge on the baseline Data.

#### Bridging performance evaluation

The bridging performance of HapBridge was evaluated using N50 and phased block N50. N50 was calculated using the WhatsHap *stats* module (see Supplementary File 2 Section 4), whereas the phased block N50 was obtained by breaking haplotype blocks at detected switch error positions and recalculating N50 to represent the contiguity of correctly phased regions. As shown in Figure 2, for ONT R9 at 80× coverage, HapCUT2-MethPhaser shows some improvement over HapCUT2 in N50 size by incorporating methylation information. However, HapCUT2-HapBridge demonstrates a substantial performance gain, increasing N50 by 142.5% over HapCUT2 and 68.6% over HapCUT2-MethPhaser, underscoring its superior ability to leverage methylation signals for long-range phasing.

**Fig. 2.**
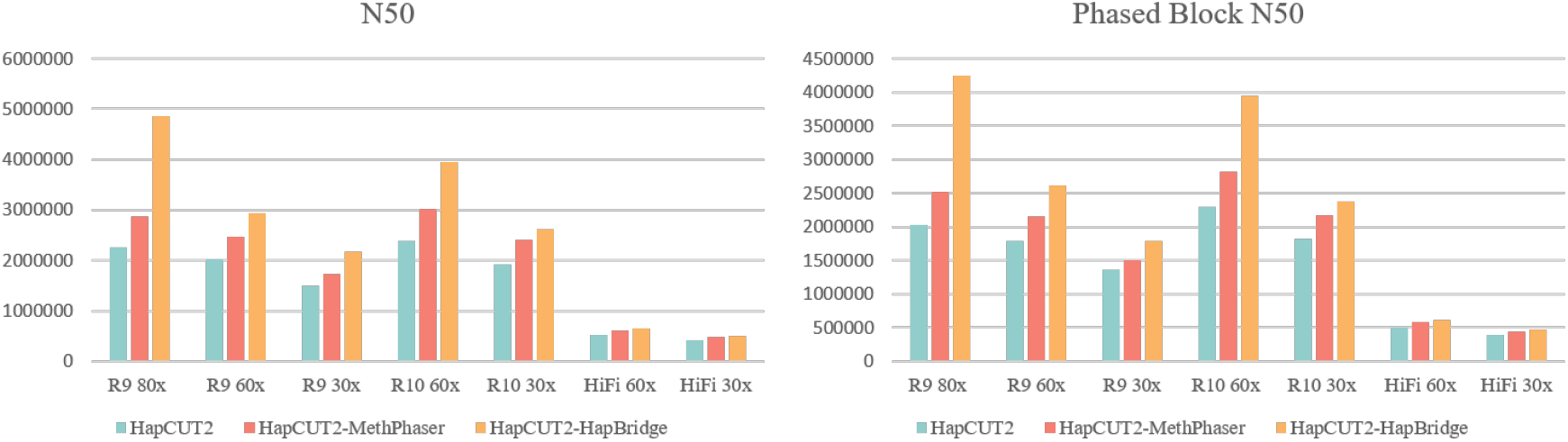
Comparison of N50 and Phased Block N50 metrics for HapCUT2, HapCUT2-MethPhaser, and HapCUT2-HapBridge on HG002 datasets including ONT R9 (80×, 60×, 30×), ONT R10 (60×, 30×), and PacBio HiFi (60×, 30×).

For ONT R10, HapCUT2-HapBridge achieves an N50 64.8% higher than HapCUT2 and 30.1% higher than HapCUT2-MethPhaser, confirming its effectiveness on improved ONT chemistries. For PacBio HiFi data, where read lengths are shorter and fewer methylation sites are available, HapCUT2-HapBridge still improves N50 by 23.6% and 6.6% compared with HapCUT2 and HapCUT2-MethPhaser, respectively, highlighting its generalizability across sequencing platforms.

The results in Table 1 show that HapCUT2-HapBridge has a higher switch error rate than HapCUT2 under low coverage conditions. Although HapCUT2-HapBridge outperforms in terms of the N50 metric, this advantage may be partially offset by connection errors within some phased blocks. To provide a more rigorous assessment of phasing accuracy while accounting for these artifacts, we further evaluated performance using the phased block N50 metric. As shown in Figure 2, HapCUT2-HapBridge consistently achieves the highest phased block N50 values across different sequencing technologies and coverage levels. For ONT R9 at 80× coverage, HapCUT2-HapBridge surpassed 4 million in phased block N50, exceeding the performance of both HapCUT2-MethPhaser and HapCUT2. At 30× coverage, HapCUT2-HapBridge maintains a competitive advantage despite the overall performance decline observed in all tools. For ONT R10 data, HapCUT2-HapBridge continues to deliver superior phased block continuity. Although shorter read lengths in PacBio HiFi data reduce N50 values across all methods, HapCUT2-HapBridge still outperforms the others. Detailed per-chromosome results for each dataset are provided in the Supplementary File 2 Tables S18-S25. Supplementary File 2 Tables S26-S34 show the results of replacing HapCUT2 with WhatsHap. Supplementary File 2 Table S38 shows the results of HG001 (60× Coverage) phasing performance comparison: WhatsHap, HapCUT2, MethPhaser, and HapBridge.

### 2.3 Ablation Experiments

To dissect the contribution of each component in HapBridge, we conducted ablation experiments by evaluating the *switch-error-correction* module and the *block-bridging* module independently across ONT R9/R10 and PacBio HiFi datasets. We benchmarked the HG002 datasets, including ONT R9 (80× and 60×), ONT R10 (60× and 30×), and HiFi (30×) sequencing data. The improvement in phasing performance was evaluated by assessing switch error and N50.

As shown in Table 3, the *switch-error-correction* module substantially reduced local phasing errors across all dataset (e.g., from 715 to 642 on ONT R9 80×), demonstrating its ability to refine haplotypes within phased blocks. In contrast, the *block-bridging* module primarily improved long-range phasing contiguity (e.g., increasing N50 from 2.26 Mb to 4.85 Mb on ONT R9 80×) while maintaining similar switch-error levels. Taken together, our results show that the two modules contribute complementary benefits: switch-error correction enhances local accuracy, whereas block bridging increases global continuity. When used jointly, HapBridge achieves the highest overall performance, indicating that methylation-guided switch detection and iterative methylation-based bridging are both necessary to maximize long-read phasing quality.

### 2.4 Assembly performance evaluation

To evaluate the downstream benefits of improved phasing, we integrated HapBridge and MethPhaser separately into the HapDup [14,32] assembly pipeline and benchmarked their performance across ONT R9/R10 and PacBio HiFi datasets(see Methods). The evaluation was conducted using three key metrics: phase block N50, intra-block switch error, and hamming error, which are generated by Merqury [27](see Supplementary File 2 Section 4).

As shown in Figure 3, compared to HapDup and HapDup-MethPhaser, HapDup-HapBridge consistently achieves significantly higher phase block N50 values across all datasets, demonstrating improved assembly continuity. For example, in the ONT R9 80× dataset, HapDup-HapBridge attains phase block N50 values of 4,423,667 (dual1) and 4,540,883 (dual2), which represent approximately a 19% increase over HapDup’s corresponding values of 3,718,884 (dual1) and 3,673,789 (dual2). It also shows a 15% improvement relative to HapDup-MethPhaser’s 4,015,109 (dual1) and 3,970,776 (dual2). These results highlight HapBridge’s ability to generate longer phased blocks and improve genome assembly continuity under various sequencing conditions.

**Fig. 3.**
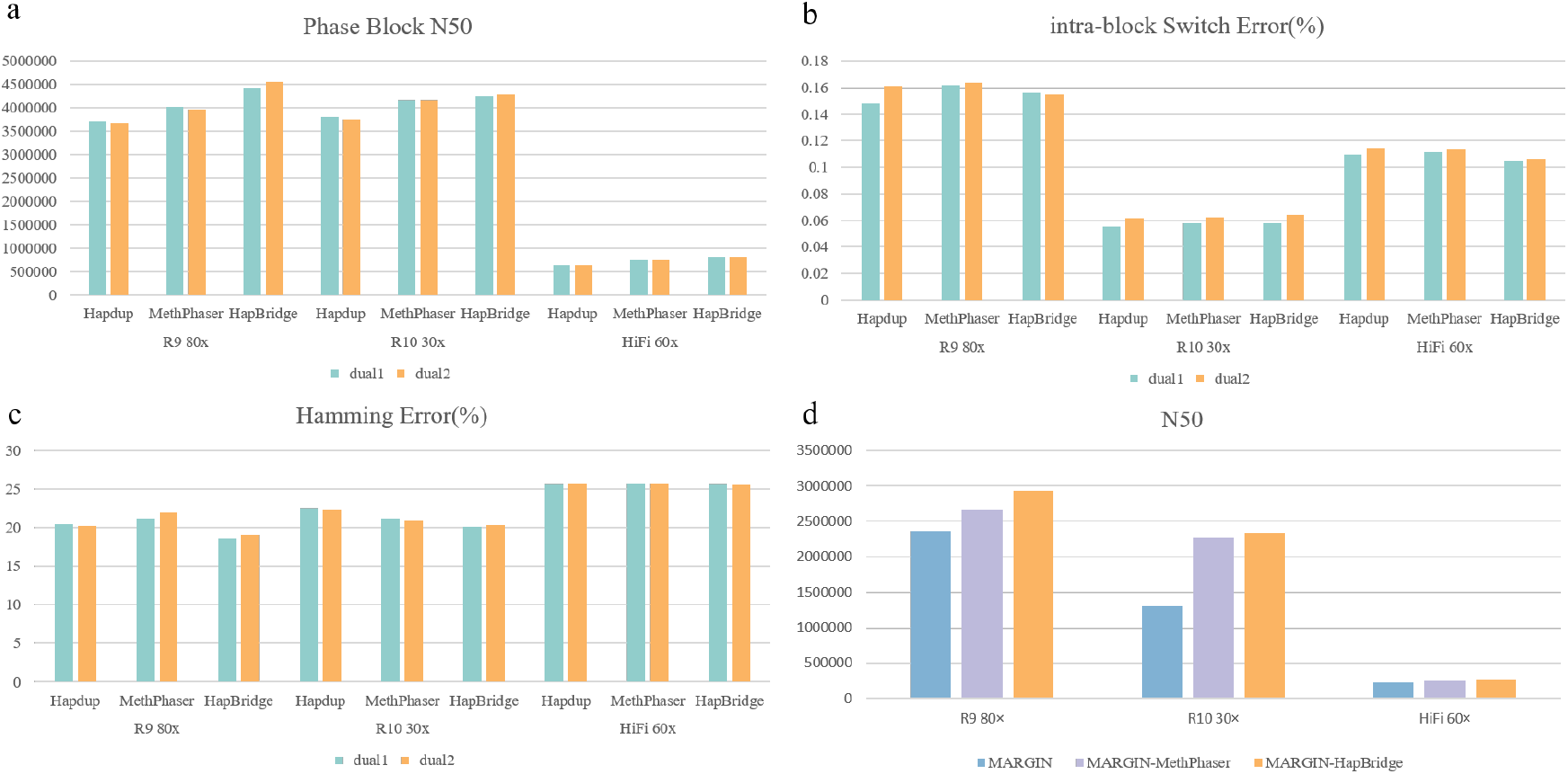
Genome assembly performance comparison across different phasing methods on the HG002 (ONT R9/R10 and HiFi) datasets. **a-c**, Phase Block N50, intra-block Switch Error (%), and Hamming Error (%) for three methods: HapDup, HapBridge, MethPhaser. **d**, Comparison of phasing results(N50) for MARGIN, MARGIN-HapBridge, and MARGIN-MethPhaser in the HapDup Pipeline We evaluated phasing tools by analyzing the N50 values of their outputs. As shown in Figure 3d, MARGIN-HapBridge achieved an N50 of 2,927,683 on ONT R9 80× data—a 24% (565,310 bp) improvement over MARGIN, HapDup’s default phasing tool. For R10 and HiFi data, N50 improved by 16% and 78%, respectively. MARGIN-MethPhaser performed intermediately between MARGIN and MARGIN-HapBridge. Additional evaluations with other SNV-based phasers combined with HapBridge are summarized in Supplementary File 2 Table S36. These results confirm that enhanced phasing improves assembly continuity, validating the effectiveness of HapBridge’s strategy.

HapDup-HapBridge achieved comparable or lower switch- and hamming-error rates than HapDup, confirming that improved contiguity does not introduce assembly artifacts. For HiFi 60× data, it achieved switch error rates of 0.104230 (dual1) and 0.106223 (dual2), outperforming HapDup (0.109354 and 0.113797). For ONT R9 80× data, its hamming error rates were 18.6206 (dual1) and 18.9672 (dual2), approximately 8% lower than HapDup’s (20.3932 and 20.2345). In contrast, HapDup-MethPhaser, despite having higher N50, showed uncompetitive error rates across all datasets. These results demonstrate that our algorithm enhances genome assembly in both continuity and accuracy.

## 3 Method

### 3.1 Calling and phasing SNPs

HapBridge starts with a Binary Alignment/Map (BAM) file, in which each read is basecalled using Guppy or Dorado, with both MM and ML tags retained to record methylation states at base resolution. These methylation tags provide the foundation for downstream haplotype-specific methylation analysis. Prior to phasing, raw reads undergo standard preprocessing, including alignment to the reference genome (using minimap2 -ax map-ont), ensuring accurate read-to-reference mapping and complete methylation annotation. Subsequently, variant calling is performed using Clair3 or Pepper, which are optimized for Oxford Nanopore and PacBio sequencing data. These tools identify single-nucleotide polymorphisms (SNPs) and generate a VCF file containing all candidate heterozygous variants. Once the VCF is obtained, phasing is performed using HapCUT2 or WhatsHap, both of which conduct SNV-based haplotype partitioning. The phased VCF is annotated with phase-set (PS) identifiers. Reads are then haplotagged using WhatsHap, assigning each read to haplotype HP1 or HP2, thereby producing a haplotagged BAM in which each read is explicitly associated with its inferred haplotype. HapBridge accepts this phased VCF and haplotagged BAM as input.

### 3.2 Detection and correction of switch errors

HapBridge next performs methylation-guided switch-error detection and correction on the phased VCF and haplotagged BAM. This method operates in two steps: (1) identification of haplotype-specific methylation sites; (2) detection of potential switch errors by calculating methylation discrepancy scores flanking each SNP.

#### Identification of haplotype-specific methylation sites

To quantitatively identify haplotype-specific methylation sites for switch error correction, HapBridge analyzes the methylation signals from haplotagged reads. For a given CpG site *i*, we collect all sequencing reads covering it. Each read *j* is associated with a haplotype tag *H*_*j*_ ∈ {HP1, HP2, Untagged} and carries a methylation value *M*_*ij*_, which is a continuous score (e.g., between 0 and 1) or a discretized state, as shown in Figure 1b, a single sequencing read contains multiple methylation signals, with each rectangle representing a unit of DNA methylation and color variations indicating differential methylation levels. Orange and cyan correspond to reads derived from distinct haplotypes, whereas yellow represents Untagged reads.

The core idea is to cluster these methylation values by haplotype and assess the distance between clusters. In Figure 1b, regions within red dashed lines show distinct haplotype-specific methylation, while those in blue dashed lines display consistent methylation across haplotypes. HapBridge utilizes haplotype-specific methylation sites to correct switch errors caused by SNP phasing tools. HapBridge collects methylation information from reads tagged with haplotype (HP) labels. We define the following for site *i*:

1. **Haplotype-Supporting Counts**: Let *S*_*i*,HP1_ and *S*_*i*,HP2_ be the sets of methylation values from reads unambiguously tagged with HP1 and HP2, respectively:

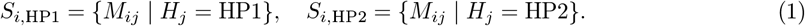

The supporting counts are the sizes of these sets: Sup_count_HP1_ = |*S*_*i*,HP1_|, Sup_count_HP2_ = |*S*_*i*,HP2_|.
2. **Haplotype-Supporting Rate**: For a haplotype (e.g., HP1), the supporting rate is the proportion of its supporting count over the total tagged reads:

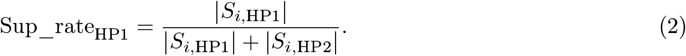
3. **Inter-Haplotype Distance**: We compute the centroid (mean methylation level) for each haplotype:

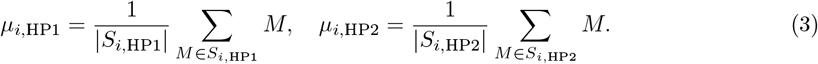

The distance between the two haplotypes at site *i* is then defined as:

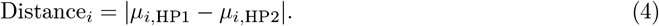

A CpG site *i* is classified as **haplotype-specific** if it satisfies all of the following conditions concurrently:

- Sup_count ≥ *th*_Sup_count_
- Sup_rate ≥ *th*_Sup_rate_
- Distance ≥ *th*_Distance_

Here, *th*_Sup_count_, *th*_Sup_rate_, and *th*_Distance_ are user-defined thresholds. The default values of 8, 0.3, and 0.3 are suitable for sequencing data with coverage greater than 60×, while 4, 0.3, and 0.3 are suitable for sequencing data with coverage less than 30×(A detailed analysis of the parameters is provided in Supplementary File 2 Section 5).

#### Localization of switch errors

Figure 1c shows the switch error caused by SNP phasing tools. Following the identification of haplotype-specific methylation sites, the next step is to determine the haplotype assignment for each individual methylation call. This is achieved through a centroid-based classification approach. Let *µ*_HP1_ and *µ*_HP2_ represent the centroid methylation values for HP1 and HP2, respectively, calculated as in Equation (3). For a methylation value *m* observed at a specific site, we compute its deviation from the midpoint between the two haplotype centroids:

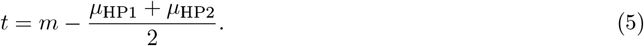

The haplotype assignment is then determined by comparing the absolute distance of *m* to each centroid:

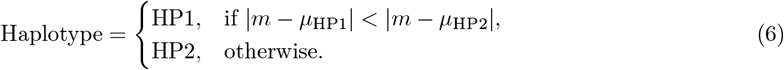

Additionally, the absolute value *t* serves as a confidence score for haplotype assignment, with larger values(we set it greater than 0.3) indicating higher confidence in the haplotype call. As shown in Figure 1d, sites are color-coded by haplotype. Without switch errors, all sites on a haplotype-assigned read should display consistent coloring. However, switch errors result in mixed patterns, as illustrated. The core premise of our detection method is that in the absence of a switch error, the haplotype-specific methylation patterns on either side of a correctly phased SNP should be consistent, supporting the same haplotype.

To quantify this effect, we introduce a **Discrepancy Score** *D* for each SNP. This score measures the change in methylation haplotype consistency when the haplotype assignments on one side of the SNP are hypothetically flipped. We identify the position with the minimum score, and if its score is below the threshold (default: –0.03), we consider it a switch error. For a given SNP *k*, the calculation proceeds as follows:

Where *c* represents the original consistency information of the methylation site, and *new*_*c* represents the consistency information after flipping.

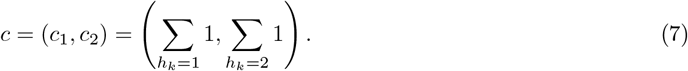

In Figure 1d, for SNP 1, *c* = (5, 6), where *c*_left_ = (2, 1) represents the haplotype distribution of the methylation site on the left, and *c*_right_ = (3, 5) represents the haplotype distribution of the methylation site on the right, flipping the haplotype of methylations on the right side of the SNP results in *new*_*c*_right_ = (5, 3).

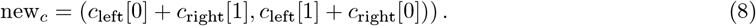

The Discrepancy Score *D*_*k*_ for SNP *k* is then defined as the normalized change in the minimum consistency:

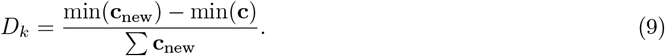

The *Discrepancy*_*Score* for SNP 1 is −0.091, and so on, for SNP 2 it is −0.273, and for SNP 3 it is −0.182. This suggests that SNP 2 is a switch error.

As validated on real sequencing data(ONT R9 80× Dataset Chromosome 22) in Figure 1d, the red data points represent the minima of the Discrepancy Score distribution, which exhibit spatial correspondence with the genomic loci identified as switch errors. The vertical axis represents the magnitude of the *Discrepancy*_*Score*, while the horizontal axis represents the position of the SNPs on the genome. The red dot marks the position 14987539, corresponding to the lowest score and indicating a switch error in the ONT R9 80x dataset.

### 3.3 Bridging two adjacent phased blocks

After correcting switch errors within phased blocks, HapBridge performs methylation-guided block bridging to connect adjacent phase blocks that lack SNV linkage(Figure 1e-f). (1) assigning haplotypes to untagged reads using haplotype-specific methylation sites; (2) refining these sites with newly tagged reads; and (3) identifying high-confidence methylation anchors between blocks via a methylation-consistency score. The process iterates until no new reads can be tagged, no further anchors are found, or bridging succeeds.

#### Haplotype assignment of untagged reads

HapBridge assigns haplotypes to untagged reads through a methylation-based voting mechanism. For each untagged read *r*, its haplotype label can be determined using the method described in Section 5.2.2. As illustrated in Figure 1e, red and blue circles represent haplotype-specific sites. For example, the third read matches the methylation distribution of *HP* 1 and is assigned accordingly, while the fourth read aligns with *HP* 2. These reads are then added to the tagged set, while untagged reads are iteratively processed for haplotype assignment.

A read is assigned to haplotype *h*\* if it meets the threshold condition:

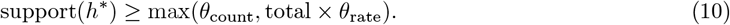

with default thresholds *θ*_count_ = 5 and *θ*_rate_ = 0.6. Finally, the tagged read is added back to the haplotagged read set to assist in determining the haplotypes of other reads until all reads have been iterated.

#### The consistency score of methylation

Methylation sites are filtered based on local signal consistency. In BAM files, methylation probabilities are encoded as values from 0 to 255, with lower values indicating less methylation. We extract and normalize methylation values from reads between phased blocks, representing low and high methylation as 0 and 1, respectively.

For any pair of methylation sites that co-occur on N reads, m of these reads exhibit a discordant methylation pattern (i.e., 01 or 10), while n reads exhibit a concordant pattern (i.e., 00 or 11), where *m* + *n* = *N*. We refer to the concordant group as being consistent at this methylation site pair and define the consistency score as follows:

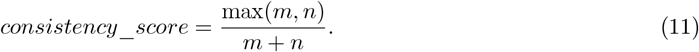

Figure 1f presents a specific example demonstrating how the consistency score is calculated. For two methylations, A and B, it is desirable for their states to predominantly occur as (0,0) and (1,1) or as (0,1) and (1,0). High consistency indicates a strong relationship between the sites. The consistency score for A and B in this example is 4/5, indicating that site A has high consistency with site B. High-consistency score methylation sites are utilized to connect phased blocks.

The algorithm computes the consistency score of each methylation site with all other sites that share several common reads. In each iteration, only the top 50% of methylation sites, ranked by consistency score, are retained. The consistency scores among the remaining sites are then recalculated, and this process is repeated iteratively until only the top 15% of methylation sites remain. These sites are considered to be reliable for downstream analysis.

#### Bridging phased blocks

Figure 1f illustrates the process of connecting two phased blocks (Phased Block 1 and Phased Block 2) using reads that span both regions, thereby generating a unified phased block. Phased Block 1(*blk*_0_) is shown on the left side of the figure, and Phased Block 2(*blk*_1_) is shown on the right, corresponding to regions defined by SNP genotyping. The intermediate reads that bridge the two blocks serve as critical links, enabling the integration of the two phased regions. These bridging reads contain multiple methylation sites, whose consistency is evaluated to assess their association with the adjacent phased blocks. Red triangles mark methylation sites with high consistency scores, indicating strong cross-read concordance and reliable phasing signals. To enhance bridging accuracy between phased blocks, the algorithm identifies haplotype-specific methylation sites during read tagging. Candidate bridging sites are selected by intersecting high-consistency sites with haplotype-specific ones (a comparison of the numbers of these two types of sites is provided in Supplementary File 2 Table S37).

Consider two adjacent phased blocks, *blk*_0_ and *blk*_1_. For all the previously filtered methylation sites, we count the number of supporting reads and record their haplotype tags (*PS, HP*). This process allows us to obtain, for each read *l*, the supported methylation locus information *sup*(*ps, hp*). If a read *l* simultaneously assigns haplotype combinations *hp*0 and *hp*1 to *blk*_0_ and *blk*_1_ respectively, we then determine whether the haplotypes are same:

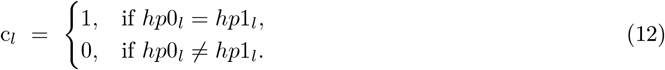

Thus, the connection support count Score(*ps*0, *ps*1, *c*) between the two blocks is incremented by one. Upon completion of read processing, the connection support counts are summarized and evaluated for compliance with the following requirements:

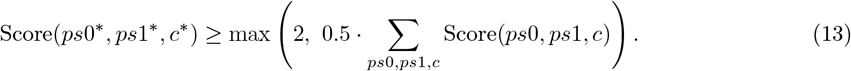

If the condition is met, (*ps*0*, *ps*1*) are declared as linked phase sets, with linkage direction:

- *c*\* = 1 indicates a direct (same haplotype) connection;
- *c*\* = −1 indicates an inverted haplotype connection;

If no triplet passes the threshold, the two blocks are considered unlinked.

### 3.4 Application of HapBridge on HapDup

To assess the utility of HapBridge in genome assembly, we integrated it into the HapDup [14,32] diploid assembly pipeline. In the original pipeline, PEPPER performs variant calling and MARGIN conducts phasing prior to haplotype-aware polishing with Flye [15]. We replaced the MARGIN with MARGIN-HapBridge for phasing and haplotagging. This configuration enables HapDup to utilize both SNV and methylation linkage information during assembly. After phasing, haplotype-tagged reads are provided to Flye for independent polishing of each haplotype, followed by structural variant resolution to finalize diploid assemblies. This integration demonstrates that HapBridge can act as a drop-in replacement for existing long-read phasers, providing enhanced methylation-guided phasing without modifying subsequent assembly steps.

## 4 Supplementary information

### 4.1 Supplementary File

**Supplementary File 1 Section 1**: Discussion; **Section 2**: Conclusions; **Section 3**: Declarations

**Supplementary File 2 Section 1**: Supplementary Tables. Tables S1-S8: Switch Error, Flip Error of HapCUT2, MethPhaser, and HapBridge on ONT R9/R10 and HiFi data (SE-Switch Error, FE-Flip Error, HE-Hamming Error). Tables S9-S17: Switch Error, Flip Error of WhatsHap, MethPhaser, and HapBridge on ONT R9/R10 and HiFi data (SE-Switch Error, FE-Flip Error, HE-Hamming Error). Tables S18-S25: The N50 and Phased Block N50 of HapCUT2, MethPhaser, and HapBridge on ONT R9/R10, and HiFi data. Tables S26-S34: The N50 and Phased Block N50 of WhatsHap, MethPhaser, and HapBridge on ONT R9/R10 and HiFi data. Table S35: The N50 values of MARGIN, HapBridge, and HapCUT2 across HiFi 60×, R9 30×, and R9 60× datasets. Table S36: Genome assembly performance comparison across different phasing methods on R9, HiFi, and R10 datasets. Block N50, Switch Error (%), and Hamming Error (%) for four methods: HapDup-MARGIN, HapDup-MARGIN(HapBridge), and HapDup-MARGIN(MethPhaser). Table S37: The comparison of the number of highly consistent methylation sites and haplotype-specific methylation sites on Chr1 in ONT R9/R10 and HiFi data. Table S38 HG001 (60× Coverage) phasing performance comparison: WhatsHap, HapCUT2, MethPhaser, and HapBridge evaluated by switch error, flip error, N50, and phased block N50

**Section 2**: Supplementary Figures. Figure S1: Flowchart of the HapBridge algorithm. Figure S2: Illustration of how HapBridge connects two phased blocks.

**Section 3**: Software Versions and Command Usage

**Section 4**: Algorithm parameters and execution commands.

**Section 5**: Pseudocode for calculating the consistency score.

### 4.2 Full Paper

A preprint can be accessed at https://doi.org/10.1101/2025.11.07.687303.

## 5 Declarations

## Acknowledgments

This work was supported in part by the National Natural Science Foundation of China under Grants (Nos. 62350004, 62332020), the Project of Xiangjiang Laboratory (No. 23XJ01011), Hunan provincial key lab on bioinformatics (2019TP1007), The high performance computing center of CSU (CSU-HighCOM01), and Fundamental Research Funds for the Central Universities of Central South University (2021zzts0208). This work was carried out in part using computing resources at the High Performance Computing Center of Central South University.

## Code availability

HapBridge is publicly available at GitHub: https://github.com/Humonex/HapBridge.

