## Supplementary File 1 for "HapBridge: A Methylation-Guided Approach for Correcting Switch Errors and Bridging Phased Blocks in Long-Read Phasing" (from another version of this preprint: version 2, 2025-11-11)

### 1. Discussion

In this study, we introduced HapBridge, a methylation-aware haplotype phasing framework that substantially improves both accuracy and continuity of long-read phasing. By intergrating allele-specific methylation signals with SNV-based haplotype information, HapBridge effectively bridges gaps between phased blocks and corrects local switch errors without the need for trio or Hi-C data. Across ONT R9/R10 and PacBio HiFi datasets, HapBridge achieved marked gains in block N50 and reduced switch errors, demonstrating its robustness across sequencing platforms and coverage levels.

Mechanistically, methylation serves as a complementary layer of linkage information that persists even in low-heterozygosity or repeat-rich regions where SNV linkage is sparse. HapBridge capitalizes on the haplotype-consistent methylation patterns to infer phase transitions between adjacent blocks. The iterative re-tagging and site-pair consistency scoring mitigate the inherent noise of methylation signals, ensuring both accuracy and stability of the bridging process. These results highlight the potential of integrating epigenetic and genetic variation to enhance the completeness of haplotype reconstruction.

We observed that HapBridge's benefit is most pronounced for ONT R10 data, reflecting improved base accuracy and methylation signal fidelity. On PacBio HiFi reads, where methylation density and read length are lower, the improvement was moderate but consistent. These findings suggest that methylation-informed phasing can adapt to diverse sequencing chemistries and could further benefit from upcoming improvements in methylation calling algorithms and raw-signal modeling.

Future work will focus on extending HapBridge to polyploid genomes and somatic mosaicism scenarios, where multiple haplotypes coexist and methylation heterogeneity increases. Additionally, integrating structural variants and transposable elements as additional linkage anchors could enable chromosome-scale phasing using long-read data alone. The demonstrated integration of HapBridge into the HapDup assembly pipeline further illustrates its practicality and potential for enhancing downstream diploid assembly and allele-specific analysis.

In summary, HapBridge establishes methylation as a reliable, sequence-independent linkage signal that enhances haplotype phasing beyond variant-based limits. By bridging the gap between genetic and epigenetic information, HapBridge offers a robust foundation for comprehensive haplotype reconstruction and downstream genomic interpretation in the long-read era.

### 2. Conclusions

We present HapBridge, a haplotype phasing method that leverages DNA methylation signals to enhance SNV-based phasing. The integration of methylation information significantly improves both phasing accuracy and block continuity across multiple sequencing platforms, including ONT and PacBio. At high coverage, HapBridge achieves the best phasing performance in terms of phased block N50, while maintaining low switch error rates. When integrated into the HapDup pipeline, HapBridge improves assembly contiguity by 8–23\%. These findings demonstrate that combining genetic and epigenetic information enables more complete and accurate haplotype reconstruction from long-read sequencing.

### 3. Declarations

- **Funding:** This work was supported in part by the National Natural Science Foundation of China under Grants (Nos. 62350004, 62332020), the Project of Xiangjiang Laboratory (No. 23XJ01011), Fundamental Research Funds for the Central Universities of Central South University (2021zzts0208).
- **Competing interests:** The authors have no competing interests to declare that are relevant to the content of this article.
- **Ethics approval and consent to participate:** Not applicable.
- **Consent for publication:** Not applicable.
- **Availability of data and materials:** The HG002 kit 10 R9 data is publicly available at https://labs.epi2me.io/gm24385-5mc-remora. the base caller is Bonito(dna_r9.4.1_e8_sup@v3.3). The original data coverage is 80x, and we used the **Samtools view** module to downsample the data to 60x and 30x. The HG002 kit 14 R10 data is publicly available at https://humanpangenome.org/data.html. The Genome in a Bottle (GIAB) SNV benchmark (V4.2.1) is publicly available at ftp://ftp-trace.ncbi.nlm.nih.gov/ReferenceSamples/giab/data/AshkenazimTrio/analysis/NIST_SVs_Integration_v0.6/. The HG001 data is publicly available at https://42basepairs.com/download/s3/ont-open-data/giab_2023.05/analysis/variant_calling/hg001_sup_60x/. The HG001 data is publicly available at: https://github.com/humanpangenomics/HG002_Data_Freeze_v1.0.
- **Code availability:** HapBridge is publicly available at GitHub https://github.com/Humonex/HapBridge
- **Author contribution:** J.W. conceived and designed this project. F.N. designed an earlier version of a workflow to phase ONT R9 data. Based on this initial workflow, H.C. designed the HapBridge method. H.C. performed experiments on ONT R9/10, and HiFi data. J.W. and F.L. provided guidance for the experiment. H.C. prepared the data and wrote the manuscript. J.W. and F.N. edited the manuscript. All authors read and approved the final manuscript.
- **Acknowledgments:** We are grateful for resources from the High-Performance Computing Center of Central South University.
