## Supplementary File 2 for "HapBridge: A Methylation-Guided Approach for Correcting Switch Errors and Bridging Phased Blocks in Long-Read Phasing" (from another version of this preprint: version 2, 2025-11-11)

### 1. Supplementary Tables

**Table S1**: Switch Error, Flip Error of HapCUT2, MethPhaser, and HapBridge on ONT R10 60× Data. (SE-Switch Error, FE- Flip Error, HE-Hamming Error)

| R10 60 | HapCUT2 | | | MethPhaser | | | HapBridge | | |
|---|---|---|---|---|---|---|---|---|---|
| | SE | FE | HE | SE | FE | HE | SE | FE | HE |
| chr1 | 56 | 22 | 1.26% | 60 | 22 | 4.68% | 53 | 22 | 0.94% |
| chr2 | 37 | 15 | 0.86% | 40 | 15 | 1.09% | 36 | 15 | 0.66% |
| chr3 | 22 | 10 | 0.44% | 22 | 10 | 0.44% | 22 | 10 | 0.37% |
| chr4 | 51 | 21 | 3.91% | 53 | 21 | 4.24% | 49 | 21 | 5.10% |
| chr5 | 29 | 10 | 2.05% | 29 | 10 | 4.60% | 28 | 10 | 5.36% |
| chr6 | 35 | 13 | 1.06% | 39 | 13 | 2.80% | 33 | 14 | 2.29% |
| chr7 | 34 | 11 | 7.24% | 36 | 11 | 9.59% | 32 | 11 | 7.20% |
| chr8 | 25 | 10 | 0.74% | 25 | 10 | 0.74% | 26 | 10 | 7.48% |
| chr9 | 30 | 13 | 0.42% | 31 | 13 | 0.61% | 30 | 13 | 2.13% |
| chr10 | 38 | 17 | 0.29% | 40 | 17 | 0.55% | 37 | 17 | 0.07% |
| chr11 | 35 | 13 | 2.55% | 37 | 13 | 5.54% | 34 | 13 | 2.00% |
| chr12 | 32 | 14 | 0.05% | 32 | 14 | 0.05% | 36 | 14 | 13.03% |
| chr13 | 13 | 5 | 1.33% | 13 | 5 | 1.37% | 13 | 5 | 1.37% |
| chr14 | 11 | 4 | 0.39% | 11 | 4 | 0.39% | 11 | 4 | 0.39% |
| chr15 | 28 | 12 | 0.45% | 29 | 12 | 2.30% | 29 | 12 | 0.46% |
| chr16 | 22 | 9 | 1.09% | 22 | 9 | 1.09% | 22 | 9 | 1.04% |
| chr17 | 28 | 13 | 0.04% | 29 | 13 | 0.34% | 28 | 13 | 0.04% |
| chr18 | 15 | 6 | 0.48% | 15 | 6 | 0.48% | 15 | 6 | 0.48% |
| chr19 | 22 | 7 | 11.83% | 22 | 7 | 20.27% | 20 | 7 | 15.70% |
| chr20 | 9 | 4 | 0.02% | 9 | 4 | 0.02% | 8 | 4 | 0.01% |
| chr21 | 20 | 9 | 2.14% | 21 | 9 | 2.85% | 20 | 9 | 2.14% |
| chr22 | 6 | 3 | 0.02% | 6 | 3 | 0.02% | 6 | 3 | 0.02% |

**Table S2** Switch Error, Flip Error of HapCUT2, MethPhaser, and HapBridge on ONT R10 30× Data (SE-Switch Error, FE- Flip Error, HE-Hamming Error)

| R10 30 | HapCUT2 | | | MethPhaser | | | HapBridge | | |
|---|---|---|---|---|---|---|---|---|---|
| | SE | FE | HE | SE | FE | HE | SE | FE | HE |
| chr1 | 58 | 22 | 2.11% | 61 | 23 | 2.62% | 59 | 22 | 2.83% |
| chr2 | 42 | 16 | 0.82% | 45 | 16 | 1.62% | 45 | 16 | 2.81% |
| chr3 | 25 | 10 | 0.42% | 27 | 10 | 1.07% | 26 | 10 | 1.28% |
| chr4 | 53 | 23 | 2.80% | 53 | 23 | 2.87% | 57 | 23 | 5.46% |
| chr5 | 26 | 9 | 2.43% | 28 | 9 | 4.88% | 27 | 9 | 2.45% |
| chr6 | 32 | 11 | 1.29% | 37 | 11 | 3.77% | 34 | 11 | 2.20% |
| chr7 | 36 | 12 | 0.57% | 39 | 12 | 0.60% | 38 | 12 | 1.41% |
| chr8 | 28 | 10 | 3.57% | 29 | 10 | 5.74% | 29 | 10 | 6.02% |
| chr9 | 32 | 13 | 4.22% | 36 | 13 | 4.67% | 33 | 13 | 5.42% |
| chr10 | 41 | 17 | 2.29% | 42 | 17 | 2.76% | 41 | 17 | 3.66% |
| chr11 | 35 | 13 | 1.68% | 37 | 13 | 3.65% | 38 | 13 | 1.93% |
| chr12 | 37 | 16 | 0.42% | 41 | 16 | 1.72% | 37 | 16 | 0.42% |
| chr13 | 19 | 7 | 2.24% | 19 | 7 | 2.24% | 20 | 7 | 2.62% |
| chr14 | 11 | 4 | 0.84% | 11 | 4 | 0.85% | 10 | 4 | 0.01% |
| chr15 | 36 | 14 | 1.35% | 38 | 14 | 1.40% | 36 | 14 | 1.36% |
| chr16 | 23 | 9 | 2.23% | 24 | 9 | 2.90% | 23 | 9 | 1.89% |
| chr17 | 31 | 13 | 0.98% | 33 | 13 | 3.51% | 32 | 13 | 1.87% |
| chr18 | 17 | 6 | 0.94% | 17 | 6 | 0.94% | 17 | 6 | 0.94% |
| chr19 | 22 | 7 | 10.86% | 22 | 7 | 10.86% | 23 | 7 | 14.11% |
| chr20 | 8 | 4 | 0.01% | 8 | 4 | 0.01% | 8 | 4 | 0.01% |
| chr21 | 21 | 9 | 2.21% | 21 | 9 | 2.21% | 24 | 9 | 24.13% |
| chr22 | 8 | 3 | 2.52% | 8 | 3 | 2.52% | 8 | 3 | 2.53% |

**Table S3** Switch Error, Flip Error of HapCUT2, MethPhaser, and HapBridge on HiFi 30× Data (SE-Switch Error, FE- Flip Error, HE-Hamming Error)

| HiFi 30 | HapCUT2 | | | MethPhaser | | | HapBridge | | |
|---|---|---|---|---|---|---|---|---|---|
| | SE | FE | HE | SE | FE | HE | SE | FE | HE |
| chr1 | 72 | 26 | 0.47% | 93 | 26 | 0.85% | 77 | 26 | 1.52% |
| chr2 | 57 | 18 | 0.54% | 78 | 18 | 1.23% | 64 | 18 | 0.97% |
| chr3 | 42 | 15 | 0.60% | 59 | 15 | 1.13% | 48 | 15 | 1.21% |
| chr4 | 62 | 24 | 0.61% | 87 | 24 | 2.35% | 63 | 24 | 0.78% |
| chr5 | 43 | 13 | 0.87% | 57 | 13 | 1.20% | 47 | 13 | 2.21% |
| chr6 | 49 | 17 | 0.35% | 73 | 18 | 0.81% | 50 | 17 | 1.21% |
| chr7 | 53 | 14 | 0.91% | 79 | 16 | 3.87% | 55 | 14 | 0.95% |
| chr8 | 41 | 14 | 0.49% | 52 | 14 | 0.72% | 47 | 14 | 2.71% |
| chr9 | 34 | 12 | 1.24% | 48 | 12 | 1.38% | 35 | 12 | 1.25% |
| chr10 | 66 | 21 | 2.92% | 72 | 21 | 3.25% | 69 | 21 | 4.88% |
| chr11 | 48 | 14 | 0.44% | 60 | 14 | 0.88% | 50 | 14 | 0.47% |
| chr12 | 48 | 16 | 2.20% | 56 | 17 | 3.19% | 49 | 16 | 2.74% |
| chr13 | 25 | 7 | 0.68% | 33 | 7 | 1.10% | 27 | 7 | 0.72% |
| chr14 | 15 | 5 | 0.55% | 22 | 5 | 1.06% | 16 | 5 | 0.56% |
| chr15 | 40 | 14 | 2.05% | 45 | 14 | 2.29% | 41 | 14 | 2.05% |
| chr16 | 30 | 9 | 0.82% | 40 | 9 | 1.96% | 35 | 9 | 1.28% |
| chr17 | 34 | 13 | 1.42% | 43 | 13 | 4.13% | 38 | 13 | 1.61% |
| chr18 | 25 | 7 | 2.02% | 31 | 7 | 3.20% | 27 | 7 | 2.03% |
| chr19 | 22 | 7 | 1.29% | 30 | 7 | 3.18% | 22 | 7 | 1.30% |
| chr20 | 12 | 4 | 0.87% | 17 | 4 | 4.76% | 13 | 4 | 1.54% |
| chr21 | 28 | 9 | 4.94% | 30 | 9 | 5.14% | 29 | 9 | 5.83% |
| chr22 | 10 | 4 | 0.16% | 17 | 4 | 2.26% | 10 | 4 | 0.16% |

**Table S4** Switch Error, Flip Error of HapCUT2, MethPhaser, and HapBridge on HiFi 60× Data (SE-Switch Error, FE- Flip Error, HE-Hamming Error)

| HiFi 60 | HapCUT2 | | | MethPhaser | | | HapBridge | | |
|---|---|---|---|---|---|---|---|---|---|
| | SE | FE | HE | SE | FE | HE | SE | FE | HE |
| chr1 | 66 | 22 | 0.75% | 77 | 22 | 1.28% | 65 | 22 | 1.34% |
| chr2 | 55 | 20 | 0.34% | 68 | 20 | 0.78% | 54 | 20 | 0.46% |
| chr3 | 40 | 10 | 0.91% | 50 | 10 | 2.02% | 40 | 10 | 1.18% |
| chr4 | 61 | 23 | 0.91% | 64 | 23 | 1.16% | 60 | 23 | 0.95% |
| chr5 | 41 | 13 | 0.69% | 47 | 13 | 0.80% | 41 | 13 | 1.15% |
| chr6 | 53 | 18 | 0.62% | 59 | 19 | 1.08% | 53 | 18 | 1.29% |
| chr7 | 51 | 12 | 1.44% | 60 | 12 | 3.40% | 50 | 12 | 1.51% |
| chr8 | 32 | 10 | 0.10% | 39 | 10 | 0.84% | 32 | 10 | 0.10% |
| chr9 | 30 | 12 | 0.04% | 34 | 12 | 0.20% | 30 | 12 | 0.60% |
| chr10 | 63 | 22 | 2.27% | 69 | 22 | 2.49% | 61 | 22 | 1.75% |
| chr11 | 50 | 14 | 1.24% | 55 | 14 | 2.05% | 50 | 14 | 1.24% |
| chr12 | 41 | 16 | 0.50% | 49 | 16 | 0.98% | 39 | 16 | 0.08% |
| chr13 | 18 | 6 | 0.61% | 23 | 6 | 1.38% | 18 | 6 | 0.61% |
| chr14 | 12 | 4 | 0.02% | 18 | 4 | 0.15% | 13 | 4 | 0.49% |
| chr15 | 34 | 13 | 0.89% | 34 | 13 | 0.89% | 34 | 13 | 0.91% |
| chr16 | 30 | 9 | 0.41% | 33 | 9 | 0.66% | 29 | 9 | 0.44% |
| chr17 | 40 | 12 | 3.06% | 47 | 12 | 3.33% | 38 | 12 | 2.93% |
| chr18 | 22 | 7 | 0.32% | 26 | 7 | 1.09% | 21 | 7 | 0.81% |
| chr19 | 22 | 7 | 3.25% | 26 | 7 | 5.38% | 22 | 7 | 5.06% |
| chr20 | 15 | 4 | 1.22% | 17 | 4 | 2.35% | 15 | 4 | 1.57% |
| chr21 | 22 | 8 | 1.61% | 24 | 8 | 4.18% | 20 | 8 | 1.10% |
| chr22 | 14 | 4 | 1.71% | 16 | 4 | 3.49% | 14 | 4 | 4.19% |

**Table S5** Switch Error, Flip Error of HapCUT2, MethPhaser, and HapBridge on ONT R9 80× Data (SE-Switch Error, FE- Flip Error, HE-Hamming Error)

| R9 80 | HapCUT2 | | | MethPhaser | | | HapBridge | | |
|---|---|---|---|---|---|---|---|---|---|
| | SE | FE | HE | SE | FE | HE | SE | FE | HE |
| chr1 | 68 | 23 | 1.99% | 68 | 23 | 2.52% | 59 | 23 | 1.41% |
| chr2 | 54 | 16 | 4.98% | 55 | 16 | 5.71% | 47 | 16 | 5.48% |
| chr3 | 34 | 10 | 1.82% | 36 | 10 | 1.92% | 27 | 10 | 3.08% |
| chr4 | 59 | 22 | 2.13% | 60 | 22 | 4.55% | 54 | 22 | 4.94% |
| chr5 | 35 | 11 | 4.89% | 39 | 11 | 5.36% | 31 | 11 | 2.09% |
| chr6 | 29 | 10 | 0.76% | 30 | 10 | 2.05% | 28 | 10 | 2.58% |
| chr7 | 41 | 11 | 3.26% | 44 | 11 | 5.27% | 42 | 11 | 3.74% |
| chr8 | 36 | 10 | 4.86% | 36 | 10 | 6.75% | 31 | 10 | 4.39% |
| chr9 | 27 | 11 | 4.30% | 29 | 11 | 5.25% | 24 | 11 | 0.63% |
| chr10 | 45 | 18 | 1.51% | 45 | 18 | 3.45% | 42 | 18 | 6.18% |
| chr11 | 43 | 14 | 2.51% | 44 | 14 | 5.28% | 38 | 13 | 2.21% |
| chr12 | 40 | 14 | 2.69% | 42 | 14 | 4.54% | 35 | 14 | 3.71% |
| chr13 | 18 | 5 | 2.94% | 18 | 5 | 5.22% | 16 | 5 | 3.23% |
| chr14 | 19 | 5 | 1.54% | 19 | 5 | 1.54% | 16 | 5 | 4.00% |
| chr15 | 34 | 14 | 0.08% | 35 | 14 | 0.11% | 33 | 14 | 0.06% |
| chr16 | 24 | 8 | 2.73% | 25 | 8 | 3.41% | 22 | 8 | 5.35% |
| chr17 | 32 | 12 | 3.52% | 33 | 12 | 8.72% | 28 | 12 | 4.69% |
| chr18 | 19 | 6 | 5.32% | 21 | 6 | 5.33% | 17 | 6 | 3.94% |
| chr19 | 21 | 7 | 4.87% | 21 | 7 | 4.87% | 19 | 7 | 23.42% |
| chr20 | 11 | 5 | 0.03% | 11 | 5 | 0.03% | 10 | 5 | 0.02% |
| chr21 | 17 | 8 | 0.05% | 17 | 8 | 0.05% | 17 | 8 | 0.05% |
| chr22 | 9 | 4 | 0.06% | 10 | 4 | 0.09% | 9 | 4 | 0.06% |

**Table S6** Switch Error, Flip Error of HapCUT2, MethPhaser, and HapBridge on NA12878 60× Data (SE-Switch Error, FE- Flip Error, HE-Hamming Error)

| NA12878 60 | HapCUT2 | | | MethPhaser | | | HapBridge | | |
|---|---|---|---|---|---|---|---|---|---|
| | SE | FE | HE | SE | FE | HE | SE | FE | HE |
| chr1 | 43 | 5 | 1.20% | 58 | 6 | 2.12% | 45 | 6 | 1.91% |
| chr2 | 68 | 13 | 0.93% | 75 | 12 | 1.88% | 73 | 14 | 1.78% |
| chr3 | 51 | 5 | 1.57% | 57 | 7 | 3.18% | 54 | 7 | 1.70% |
| chr4 | 41 | 8 | 1.13% | 54 | 9 | 1.67% | 42 | 9 | 1.32% |
| chr5 | 29 | 3 | 0.83% | 39 | 3 | 2.61% | 33 | 4 | 1.09% |
| chr6 | 37 | 4 | 1.40% | 43 | 4 | 2.26% | 38 | 4 | 1.50% |
| chr7 | 31 | 4 | 0.71% | 38 | 5 | 2.56% | 34 | 5 | 1.27% |
| chr8 | 38 | 7 | 1.38% | 43 | 9 | 2.59% | 39 | 7 | 2.70% |
| chr9 | 24 | 4 | 2.19% | 30 | 5 | 2.46% | 26 | 4 | 2.96% |
| chr10 | 22 | 2 | 0.73% | 26 | 2 | 0.90% | 27 | 4 | 4.31% |
| chr11 | 26 | 5 | 0.87% | 28 | 6 | 1.45% | 31 | 7 | 1.35% |
| chr12 | 28 | 5 | 1.34% | 36 | 8 | 1.37% | 33 | 8 | 1.47% |
| chr13 | 24 | 9 | 0.03% | 28 | 9 | 0.83% | 25 | 10 | 0.03% |
| chr14 | 19 | 2 | 0.42% | 25 | 3 | 1.96% | 19 | 3 | 0.75% |
| chr15 | 18 | 4 | 0.96% | 22 | 4 | 2.64% | 18 | 4 | 1.01% |
| chr16 | 21 | 6 | 2.08% | 22 | 7 | 2.08% | 23 | 6 | 6.96% |
| chr17 | 17 | 1 | 0.81% | 20 | 2 | 1.38% | 20 | 5 | 1.31% |
| chr18 | 15 | 1 | 0.49% | 18 | 1 | 0.56% | 18 | 2 | 2.10% |
| chr19 | 15 | 1 | 2.30% | 17 | 1 | 3.87% | 16 | 2 | 2.49% |
| chr20 | 16 | 2 | 0.61% | 19 | 2 | 5.16% | 14 | 2 | 0.87% |
| chr21 | 6 | 2 | 1.61% | 6 | 2 | 1.61% | 6 | 2 | 2.39% |
| chr22 | 7 | 1 | 1.45% | 8 | 1 | 6.81% | 8 | 1 | 0.76% |

**Table S7** Switch Error, Flip Error of HapCUT2, MethPhaser, and HapBridge on ONT R9 30× Data (SE-Switch Error, FE- Flip Error, HE-Hamming Error)

| R9 30 | HapCUT2 | | | MethPhaser | | | HapBridge | | |
|---|---|---|---|---|---|---|---|---|---|
| | SE | FE | HE | SE | FE | HE | SE | FE | HE |
| chr1 | 79 | 23 | 4.34% | 89 | 23 | 7.47% | 80 | 23 | 4.69% |
| chr2 | 68 | 17 | 3.13% | 74 | 18 | 4.05% | 68 | 17 | 3.48% |
| chr3 | 55 | 12 | 5.29% | 58 | 12 | 6.80% | 55 | 12 | 5.29% |
| chr4 | 69 | 20 | 5.17% | 76 | 20 | 7.51% | 69 | 20 | 5.17% |
| chr5 | 55 | 11 | 3.57% | 61 | 11 | 4.19% | 55 | 11 | 4.32% |
| chr6 | 46 | 10 | 2.54% | 48 | 10 | 3.90% | 46 | 10 | 2.54% |
| chr7 | 62 | 13 | 6.19% | 68 | 13 | 6.49% | 62 | 13 | 6.19% |
| chr8 | 39 | 9 | 5.28% | 41 | 9 | 5.72% | 39 | 9 | 5.28% |
| chr9 | 38 | 12 | 7.64% | 40 | 12 | 7.96% | 38 | 12 | 7.64% |
| chr10 | 57 | 16 | 4.60% | 58 | 17 | 4.60% | 58 | 16 | 4.65% |
| chr11 | 59 | 14 | 5.09% | 64 | 14 | 6.04% | 59 | 14 | 5.14% |
| chr12 | 46 | 16 | 3.11% | 52 | 16 | 4.82% | 46 | 16 | 3.11% |
| chr13 | 20 | 4 | 4.71% | 22 | 4 | 5.47% | 20 | 4 | 4.71% |
| chr14 | 20 | 4 | 2.07% | 22 | 4 | 3.71% | 20 | 4 | 2.07% |
| chr15 | 40 | 13 | 6.46% | 40 | 13 | 6.47% | 39 | 13 | 6.44% |
| chr16 | 28 | 8 | 1.73% | 31 | 8 | 2.23% | 28 | 8 | 1.85% |
| chr17 | 38 | 10 | 3.65% | 38 | 10 | 3.65% | 38 | 10 | 3.65% |
| chr18 | 28 | 6 | 3.58% | 29 | 6 | 4.79% | 28 | 6 | 3.58% |
| chr19 | 18 | 6 | 0.09% | 18 | 6 | 4.65% | 18 | 6 | 0.09% |
| chr20 | 14 | 4 | 2.65% | 14 | 4 | 5.59% | 14 | 4 | 2.65% |
| chr21 | 17 | 7 | 2.10% | 19 | 7 | 5.66% | 17 | 7 | 2.10% |
| chr22 | 14 | 4 | 5.79% | 14 | 4 | 5.79% | 14 | 4 | 5.79% |

**Table S8** Switch Error, Flip Error of HapCUT2, MethPhaser, and HapBridge on ONT R9 60× Data (SE-Switch Error, FE- Flip Error, HE-Hamming Error)

| R9 60 | HapCUT2 | | | MethPhaser | | | HapBridge | | |
|---|---|---|---|---|---|---|---|---|---|
| | SE | FE | HE | SE | FE | HE | SE | FE | HE |
| chr1 | 76 | 21 | 5.46% | 77 | 21 | 5.92% | 65 | 21 | 3.68% |
| chr2 | 50 | 15 | 3.36% | 51 | 15 | 4.29% | 48 | 15 | 3.76% |
| chr3 | 54 | 13 | 7.08% | 56 | 13 | 9.33% | 47 | 12 | 9.34% |
| chr4 | 58 | 20 | 4.87% | 60 | 20 | 8.08% | 53 | 20 | 4.36% |
| chr5 | 40 | 10 | 6.96% | 41 | 10 | 7.13% | 36 | 10 | 4.43% |
| chr6 | 36 | 12 | 0.37% | 38 | 12 | 0.86% | 34 | 11 | 0.37% |
| chr7 | 45 | 11 | 8.04% | 46 | 11 | 10.07% | 43 | 11 | 8.22% |
| chr8 | 36 | 10 | 4.97% | 37 | 10 | 5.14% | 30 | 10 | 3.27% |
| chr9 | 24 | 11 | 1.10% | 25 | 11 | 1.10% | 22 | 11 | 0.02% |
| chr10 | 47 | 17 | 3.65% | 48 | 17 | 3.73% | 43 | 18 | 2.77% |
| chr11 | 49 | 16 | 3.26% | 49 | 16 | 4.18% | 46 | 16 | 4.81% |
| chr12 | 46 | 15 | 5.18% | 47 | 15 | 10.11% | 41 | 15 | 3.34% |
| chr13 | 18 | 5 | 5.42% | 18 | 5 | 5.69% | 17 | 5 | 5.35% |
| chr14 | 13 | 4 | 0.58% | 13 | 4 | 0.58% | 14 | 4 | 1.66% |
| chr15 | 39 | 14 | 2.48% | 42 | 14 | 3.71% | 36 | 14 | 2.11% |
| chr16 | 28 | 9 | 1.30% | 29 | 9 | 4.35% | 26 | 9 | 3.31% |
| chr17 | 30 | 12 | 0.79% | 30 | 12 | 0.79% | 28 | 12 | 0.57% |
| chr18 | 22 | 7 | 6.88% | 22 | 7 | 6.88% | 22 | 7 | 6.88% |
| chr19 | 23 | 7 | 8.71% | 24 | 7 | 9.19% | 20 | 7 | 13.44% |
| chr20 | 14 | 4 | 3.97% | 14 | 4 | 3.97% | 11 | 4 | 3.96% |
| chr21 | 17 | 8 | 1.49% | 17 | 8 | 1.49% | 17 | 8 | 1.49% |
| chr22 | 7 | 3 | 0.04% | 7 | 3 | 0.04% | 7 | 3 | 0.04% |

**Table S9** Switch Error, Flip Error of WhasHap, MethPhaser, and HapBridge on the baseline Data.(SE-Switch Error, FE- Flip Error, HE-Hamming Error)

| | WhasHap | | MethPhaser | | HapBridge | |
|---|---|---|---|---|---|---|
| data | SE | FE | SE | FE | SE | FE |
| R9 80× | 717 | 253 | 758 | 254 | 658 | 255 |
| R9 60× | 774 | 252 | 807 | 255 | 741 | 253 |
| R9 30× | 939 | 239 | 1005 | 241 | 974 | 240 |
| R10 60× | 6132 | 2890 | 6142 | 2891 | 6061 | 2891 |
| R10 30× | 618 | 232 | 657 | 233 | 664 | 232 |
| HiFi 60× | 713 | 238 | 827 | 239 | 715 | 239 |
| HiFi 30× | 756 | 240 | 1039 | 245 | 812 | 240 |

**Table S10** Switch Error, Flip Error of WhasHap, MethPhaser, and HapBridge on NA12878 60× Data (SE-Switch Error, FE- Flip Error, HE-Hamming Error)

| NA12878 60 | WhasHap | | | MethPhaser | | | HapBridge | | |
|---|---|---|---|---|---|---|---|---|---|
| | SE | FE | HE | SE | FE | HE | SE | FE | HE |
| chr1 | 44 | 5 | 0.92% | 62 | 6 | 2.05% | 55 | 10 | 1.92% |
| chr2 | 64 | 12 | 1.70% | 73 | 14 | 2.87% | 69 | 13 | 1.70% |
| chr3 | 49 | 3 | 1.93% | 59 | 5 | 4.66% | 57 | 8 | 1.56% |
| chr4 | 31 | 2 | 1.58% | 43 | 4 | 2.46% | 40 | 6 | 1.93% |
| chr5 | 27 | 1 | 0.90% | 34 | 1 | 2.86% | 31 | 2 | 1.01% |
| chr6 | 32 | 2 | 1.42% | 41 | 3 | 2.79% | 36 | 2 | 2.86% |
| chr7 | 28 | 3 | 1.23% | 32 | 4 | 2.84% | 31 | 5 | 2.38% |
| chr8 | 29 | 6 | 1.21% | 33 | 7 | 1.84% | 32 | 7 | 0.84% |
| chr9 | 21 | 5 | 1.48% | 26 | 6 | 1.67% | 27 | 6 | 1.96% |
| chr10 | 21 | 1 | 1.38% | 26 | 1 | 2.20% | 26 | 4 | 3.76% |
| chr11 | 23 | 4 | 0.99% | 27 | 6 | 1.69% | 25 | 6 | 1.25% |
| chr12 | 27 | 4 | 0.25% | 30 | 5 | 0.26% | 34 | 7 | 0.61% |
| chr13 | 11 | 1 | 1.00% | 15 | 1 | 3.36% | 13 | 2 | 1.24% |
| chr14 | 13 | 0 | 1.15% | 17 | 1 | 3.23% | 17 | 2 | 4.64% |
| chr15 | 12 | 3 | 0.03% | 17 | 4 | 0.25% | 17 | 3 | 0.65% |
| chr16 | 18 | 4 | 0.88% | 18 | 4 | 0.88% | 19 | 4 | 4.31% |
| chr17 | 20 | 2 | 0.19% | 24 | 3 | 0.21% | 25 | 6 | 1.60% |
| chr18 | 11 | 0 | 0.48% | 15 | 0 | 0.71% | 13 | 1 | 0.48% |
| chr19 | 9 | 0 | 2.16% | 11 | 1 | 2.84% | 8 | 1 | 1.34% |
| chr20 | 10 | 2 | 0.03% | 11 | 2 | 0.56% | 12 | 2 | 1.02% |
| chr21 | 6 | 2 | 1.61% | 6 | 2 | 1.61% | 7 | 2 | 2.39% |
| chr22 | 8 | 1 | 1.43% | 10 | 1 | 6.76% | 12 | 3 | 1.43% |

**Table S11** Switch Error, Flip Error of WhasHap, MethPhaser, and HapBridge on ONT R9 60× Data (SE-Switch Error, FE- Flip Error, HE-Hamming Error)

| R9 60 | WhasHap | | | MethPhaser | | | HapBridge | | |
|---|---|---|---|---|---|---|---|---|---|
| | SE | FE | HE | SE | FE | HE | SE | FE | HE |
| chr1 | 70 | 21 | 3.98% | 75 | 22 | 9.98% | 65 | 21 | 4.43% |
| chr2 | 54 | 16 | 4.11% | 54 | 16 | 4.99% | 53 | 16 | 5.55% |
| chr3 | 43 | 10 | 5.45% | 45 | 11 | 7.78% | 42 | 11 | 7.04% |
| chr4 | 58 | 22 | 2.76% | 59 | 22 | 8.06% | 57 | 22 | 3.19% |
| chr5 | 41 | 11 | 3.74% | 44 | 11 | 4.41% | 41 | 11 | 4.41% |
| chr6 | 33 | 11 | 0.88% | 35 | 11 | 1.35% | 31 | 11 | 1.23% |
| chr7 | 42 | 13 | 2.42% | 43 | 13 | 3.08% | 42 | 13 | 2.86% |
| chr8 | 36 | 9 | 5.97% | 38 | 9 | 6.98% | 31 | 9 | 7.83% |
| chr9 | 32 | 14 | 0.76% | 34 | 14 | 3.53% | 31 | 14 | 1.81% |
| chr10 | 49 | 17 | 3.25% | 49 | 17 | 5.23% | 47 | 17 | 2.51% |
| chr11 | 49 | 15 | 3.38% | 52 | 15 | 5.32% | 42 | 15 | 3.56% |
| chr12 | 44 | 15 | 1.41% | 46 | 16 | 4.73% | 44 | 15 | 5.06% |
| chr13 | 26 | 7 | 7.13% | 29 | 7 | 8.83% | 23 | 7 | 7.67% |
| chr14 | 17 | 5 | 1.16% | 17 | 5 | 1.16% | 18 | 5 | 1.75% |
| chr15 | 38 | 15 | 3.46% | 40 | 15 | 4.91% | 37 | 15 | 1.63% |
| chr16 | 29 | 10 | 0.19% | 29 | 10 | 1.16% | 28 | 10 | 1.21% |
| chr17 | 37 | 13 | 2.00% | 38 | 13 | 2.57% | 36 | 13 | 1.43% |
| chr18 | 22 | 6 | 5.93% | 24 | 6 | 6.85% | 20 | 6 | 5.78% |
| chr19 | 16 | 6 | 1.75% | 17 | 6 | 6.51% | 16 | 6 | 6.33% |
| chr20 | 14 | 5 | 3.46% | 14 | 5 | 4.36% | 13 | 5 | 3.37% |
| chr21 | 16 | 8 | 0.04% | 16 | 8 | 0.04% | 16 | 8 | 0.04% |
| chr22 | 8 | 3 | 0.06% | 9 | 3 | 0.08% | 8 | 3 | 2.60% |

**Table S12** Switch Error, Flip Error of WhasHap, MethPhaser, and HapBridge on HiFi 60× Data (SE-Switch Error, FE- Flip Error, HE-Hamming Error)

| HiFi 60 | WhasHap | | | MethPhaser | | | HapBridge | | |
|---|---|---|---|---|---|---|---|---|---|
| | SE | FE | HE | SE | FE | HE | SE | FE | HE |
| chr1 | 60 | 21 | 0.37% | 71 | 21 | 1.34% | 61 | 21 | 0.42% |
| chr2 | 51 | 17 | 0.35% | 60 | 17 | 0.97% | 50 | 17 | 0.52% |
| chr3 | 34 | 11 | 0.76% | 39 | 11 | 0.97% | 34 | 11 | 1.28% |
| chr4 | 48 | 21 | 0.86% | 49 | 21 | 0.87% | 48 | 21 | 0.89% |
| chr5 | 36 | 10 | 1.44% | 43 | 10 | 2.40% | 36 | 10 | 1.46% |
| chr6 | 37 | 11 | 0.59% | 47 | 12 | 1.04% | 38 | 11 | 1.27% |
| chr7 | 43 | 11 | 1.43% | 49 | 11 | 1.83% | 41 | 11 | 2.07% |
| chr8 | 33 | 10 | 0.92% | 40 | 10 | 1.21% | 32 | 10 | 0.14% |
| chr9 | 26 | 12 | 0.07% | 29 | 12 | 0.21% | 28 | 12 | 0.11% |
| chr10 | 48 | 16 | 1.03% | 52 | 16 | 2.32% | 49 | 17 | 1.72% |
| chr11 | 42 | 12 | 1.80% | 46 | 12 | 1.93% | 42 | 12 | 2.13% |
| chr12 | 37 | 15 | 0.06% | 42 | 15 | 0.39% | 38 | 15 | 0.07% |
| chr13 | 20 | 5 | 0.71% | 25 | 5 | 1.63% | 19 | 5 | 0.69% |
| chr14 | 14 | 5 | 0.06% | 20 | 5 | 1.10% | 15 | 5 | 0.54% |
| chr15 | 31 | 12 | 0.36% | 34 | 12 | 0.65% | 31 | 12 | 0.13% |
| chr16 | 27 | 9 | 0.49% | 30 | 9 | 1.08% | 26 | 9 | 0.53% |
| chr17 | 38 | 12 | 2.06% | 51 | 12 | 3.16% | 38 | 12 | 3.83% |
| chr18 | 19 | 6 | 0.31% | 20 | 6 | 0.33% | 19 | 6 | 0.80% |
| chr19 | 20 | 6 | 2.92% | 23 | 6 | 4.43% | 20 | 6 | 5.29% |
| chr20 | 15 | 4 | 1.02% | 19 | 4 | 1.61% | 16 | 4 | 1.99% |
| chr21 | 21 | 8 | 2.09% | 23 | 8 | 2.22% | 20 | 8 | 3.03% |
| chr22 | 13 | 4 | 3.55% | 15 | 4 | 4.42% | 14 | 4 | 6.15% |

**Table S13** Switch Error, Flip Error of WhasHap, MethPhaser, and HapBridge on HiFi 30× Data (SE-Switch Error, FE- Flip Error, HE-Hamming Error)

| HiFi 30 | WhasHap | | | MethPhaser | | | HapBridge | | |
|---|---|---|---|---|---|---|---|---|---|
| | SE | FE | HE | SE | FE | HE | SE | FE | HE |
| chr1 | 61 | 22 | 0.39% | 85 | 22 | 1.33% | 67 | 22 | 1.85% |
| chr2 | 55 | 17 | 0.52% | 81 | 17 | 1.67% | 62 | 17 | 1.15% |
| chr3 | 34 | 12 | 0.72% | 55 | 12 | 1.47% | 42 | 12 | 1.64% |
| chr4 | 55 | 21 | 0.10% | 76 | 21 | 1.34% | 55 | 21 | 0.14% |
| chr5 | 39 | 10 | 0.76% | 57 | 10 | 0.98% | 43 | 10 | 0.88% |
| chr6 | 38 | 11 | 0.33% | 53 | 11 | 3.54% | 39 | 11 | 0.39% |
| chr7 | 47 | 12 | 0.95% | 70 | 14 | 3.96% | 49 | 12 | 1.09% |
| chr8 | 32 | 10 | 0.52% | 43 | 10 | 0.69% | 37 | 10 | 2.75% |
| chr9 | 33 | 12 | 1.52% | 48 | 13 | 2.02% | 37 | 12 | 1.62% |
| chr10 | 54 | 16 | 1.61% | 62 | 16 | 2.95% | 56 | 16 | 2.28% |
| chr11 | 44 | 12 | 1.21% | 63 | 14 | 1.84% | 46 | 12 | 1.56% |
| chr12 | 41 | 15 | 1.16% | 51 | 15 | 3.02% | 42 | 15 | 2.30% |
| chr13 | 17 | 5 | 1.06% | 33 | 5 | 1.60% | 20 | 5 | 1.14% |
| chr14 | 14 | 5 | 0.56% | 21 | 5 | 1.06% | 14 | 5 | 0.56% |
| chr15 | 36 | 12 | 1.79% | 40 | 12 | 2.14% | 37 | 12 | 1.80% |
| chr16 | 27 | 8 | 1.07% | 37 | 8 | 3.35% | 32 | 8 | 1.95% |
| chr17 | 38 | 12 | 1.10% | 48 | 12 | 2.79% | 39 | 12 | 2.07% |
| chr18 | 20 | 6 | 0.57% | 26 | 6 | 1.60% | 23 | 6 | 0.59% |
| chr19 | 23 | 6 | 2.45% | 32 | 6 | 3.94% | 24 | 6 | 4.15% |
| chr20 | 14 | 4 | 0.44% | 20 | 4 | 3.54% | 14 | 4 | 0.45% |
| chr21 | 22 | 8 | 2.27% | 25 | 8 | 2.34% | 22 | 8 | 3.21% |
| chr22 | 12 | 4 | 0.43% | 13 | 4 | 0.61% | 12 | 4 | 1.84% |

**Table S14** Switch Error, Flip Error of WhasHap, MethPhaser, and HapBridge on ONT R9 80× Data (SE-Switch Error, FE- Flip Error, HE-Hamming Error)

| R9 80 | WhasHap | | | MethPhaser | | | HapBridge | | |
|---|---|---|---|---|---|---|---|---|---|
| | SE | FE | HE | SE | FE | HE | SE | FE | HE |
| chr1 | 67 | 23 | 1.91% | 69 | 24 | 1.91% | 60 | 23 | 2.30% |
| chr2 | 55 | 19 | 3.85% | 60 | 19 | 5.91% | 51 | 19 | 3.85% |
| chr3 | 24 | 9 | 0.50% | 26 | 9 | 1.30% | 24 | 9 | 0.80% |
| chr4 | 55 | 21 | 2.36% | 60 | 21 | 6.05% | 48 | 21 | 1.09% |
| chr5 | 34 | 9 | 3.73% | 36 | 9 | 5.07% | 32 | 9 | 3.20% |
| chr6 | 39 | 14 | 0.75% | 42 | 14 | 2.40% | 38 | 14 | 1.71% |
| chr7 | 42 | 13 | 6.33% | 45 | 13 | 7.55% | 40 | 13 | 1.94% |
| chr8 | 32 | 9 | 3.44% | 33 | 9 | 6.11% | 31 | 10 | 6.77% |
| chr9 | 25 | 11 | 2.13% | 27 | 11 | 2.14% | 24 | 11 | 1.08% |
| chr10 | 46 | 18 | 1.52% | 47 | 18 | 2.14% | 43 | 18 | 0.81% |
| chr11 | 49 | 16 | 0.88% | 52 | 16 | 1.59% | 42 | 16 | 0.62% |
| chr12 | 40 | 15 | 0.43% | 43 | 15 | 3.42% | 40 | 16 | 1.16% |
| chr13 | 24 | 7 | 6.95% | 27 | 7 | 8.10% | 20 | 7 | 3.50% |
| chr14 | 17 | 5 | 1.51% | 17 | 5 | 1.51% | 15 | 5 | 2.32% |
| chr15 | 30 | 13 | 1.65% | 31 | 13 | 1.70% | 29 | 13 | 1.62% |
| chr16 | 24 | 9 | 0.35% | 25 | 9 | 0.43% | 22 | 9 | 1.26% |
| chr17 | 36 | 13 | 2.13% | 37 | 13 | 2.83% | 30 | 13 | 0.22% |
| chr18 | 21 | 6 | 4.50% | 22 | 6 | 5.46% | 17 | 6 | 4.74% |
| chr19 | 16 | 6 | 5.08% | 17 | 6 | 10.23% | 15 | 6 | 8.99% |
| chr20 | 12 | 5 | 0.03% | 12 | 5 | 0.03% | 10 | 5 | 0.02% |
| chr21 | 19 | 8 | 0.41% | 19 | 8 | 0.41% | 18 | 8 | 0.40% |
| chr22 | 10 | 4 | 0.08% | 11 | 4 | 4.02% | 9 | 4 | 0.07% |

**Table S15** Switch Error, Flip Error of WhasHap, MethPhaser, and HapBridge on ONT R10 60× Data (SE-Switch Error, FE- Flip Error, HE-Hamming Error)

| R10 60 | WhasHap | | | MethPhaser | | | HapBridge | | |
|---|---|---|---|---|---|---|---|---|---|
| | SE | FE | HE | SE | FE | HE | SE | FE | HE |
| chr1 | 522 | 247 | 3.78% | 523 | 247 | 6.91% | 519 | 247 | 2.38% |
| chr2 | 566 | 265 | 8.42% | 568 | 266 | 8.70% | 562 | 267 | 7.39% |
| chr3 | 442 | 217 | 4.03% | 444 | 217 | 5.09% | 440 | 218 | 0.92% |
| chr4 | 404 | 191 | 9.53% | 405 | 191 | 10.68% | 402 | 191 | 12.11% |
| chr5 | 365 | 170 | 12.52% | 368 | 170 | 17.10% | 358 | 170 | 13.38% |
| chr6 | 382 | 184 | 3.44% | 382 | 184 | 5.30% | 376 | 183 | 2.57% |
| chr7 | 402 | 186 | 11.35% | 402 | 186 | 13.14% | 397 | 185 | 11.41% |
| chr8 | 315 | 146 | 10.11% | 315 | 146 | 13.57% | 311 | 147 | 17.27% |
| chr9 | 280 | 131 | 6.82% | 280 | 131 | 9.97% | 274 | 131 | 2.96% |
| chr10 | 302 | 145 | 7.44% | 302 | 145 | 7.52% | 301 | 145 | 6.89% |
| chr11 | 220 | 98 | 13.07% | 220 | 98 | 14.01% | 213 | 98 | 12.25% |
| chr12 | 322 | 153 | 10.67% | 323 | 153 | 14.08% | 317 | 154 | 6.38% |
| chr13 | 157 | 73 | 2.37% | 157 | 73 | 2.72% | 157 | 73 | 2.65% |
| chr14 | 189 | 91 | 7.39% | 189 | 91 | 23.18% | 185 | 90 | 5.12% |
| chr15 | 185 | 86 | 6.81% | 186 | 86 | 13.70% | 184 | 86 | 10.23% |
| chr16 | 227 | 109 | 7.37% | 227 | 109 | 9.28% | 227 | 109 | 16.06% |
| chr17 | 213 | 97 | 14.06% | 213 | 97 | 16.46% | 211 | 97 | 2.90% |
| chr18 | 172 | 79 | 18.55% | 171 | 79 | 19.02% | 169 | 79 | 19.59% |
| chr19 | 218 | 104 | 9.12% | 218 | 104 | 9.12% | 215 | 104 | 21.02% |
| chr20 | 89 | 42 | 10.00% | 89 | 42 | 10.00% | 85 | 42 | 0.36% |
| chr21 | 79 | 36 | 6.01% | 79 | 36 | 6.01% | 78 | 36 | 24.73% |
| chr22 | 81 | 40 | 1.74% | 81 | 40 | 1.74% | 80 | 39 | 14.62% |

**Table S16** Switch Error, Flip Error of WhasHap, MethPhaser, and HapBridge on ONT R10 30× Data (SE-Switch Error, FE- Flip Error, HE-Hamming Error)

| R10 30 | WhasHap | | | MethPhaser | | | HapBridge | | |
|---|---|---|---|---|---|---|---|---|---|
| | SE | FE | HE | SE | FE | HE | SE | FE | HE |
| chr1 | 51 | 21 | 0.84% | 55 | 22 | 1.35% | 51 | 21 | 1.35% |
| chr2 | 44 | 17 | 1.63% | 48 | 17 | 2.28% | 52 | 17 | 3.46% |
| chr3 | 25 | 10 | 1.28% | 28 | 10 | 4.26% | 31 | 10 | 2.76% |
| chr4 | 50 | 21 | 4.17% | 53 | 21 | 5.25% | 53 | 21 | 3.42% |
| chr5 | 28 | 10 | 1.58% | 29 | 10 | 1.59% | 32 | 10 | 7.07% |
| chr6 | 29 | 10 | 0.52% | 31 | 10 | 0.54% | 32 | 10 | 2.49% |
| chr7 | 38 | 11 | 2.58% | 40 | 11 | 4.27% | 41 | 11 | 9.60% |
| chr8 | 29 | 10 | 3.20% | 31 | 10 | 3.21% | 31 | 10 | 12.16% |
| chr9 | 26 | 12 | 0.51% | 28 | 12 | 0.59% | 26 | 12 | 0.58% |
| chr10 | 37 | 15 | 2.36% | 38 | 15 | 3.75% | 37 | 15 | 3.20% |
| chr11 | 34 | 12 | 1.35% | 35 | 12 | 3.05% | 37 | 12 | 3.61% |
| chr12 | 39 | 15 | 5.61% | 42 | 15 | 10.20% | 38 | 15 | 4.54% |
| chr13 | 14 | 5 | 0.96% | 17 | 5 | 1.00% | 17 | 5 | 1.95% |
| chr14 | 12 | 4 | 0.88% | 12 | 4 | 2.93% | 15 | 4 | 5.23% |
| chr15 | 34 | 12 | 1.74% | 37 | 12 | 6.12% | 33 | 12 | 1.34% |
| chr16 | 22 | 9 | 2.22% | 24 | 9 | 2.90% | 25 | 9 | 5.55% |
| chr17 | 29 | 11 | 1.08% | 30 | 11 | 3.93% | 29 | 11 | 1.12% |
| chr18 | 20 | 6 | 1.81% | 21 | 6 | 1.82% | 23 | 6 | 2.81% |
| chr19 | 21 | 6 | 15.08% | 22 | 6 | 19.89% | 20 | 6 | 18.90% |
| chr20 | 8 | 4 | 0.01% | 8 | 4 | 0.01% | 10 | 4 | 0.62% |
| chr21 | 20 | 8 | 2.25% | 20 | 8 | 2.25% | 22 | 8 | 20.35% |
| chr22 | 8 | 3 | 2.52% | 8 | 3 | 2.52% | 9 | 3 | 2.53% |

**Table S17** Switch Error, Flip Error of WhasHap, MethPhaser, and HapBridge on ONT R9 30× Data (SE-Switch Error, FE- Flip Error, HE-Hamming Error)

| R9 30 | WhasHap | | | MethPhaser | | | HapBridge | | |
|---|---|---|---|---|---|---|---|---|---|
| | SE | FE | HE | SE | FE | HE | SE | FE | HE |
| chr1 | 73 | 20 | 5.17% | 82 | 20 | 8.44% | 74 | 20 | 5.92% |
| chr2 | 74 | 17 | 2.71% | 78 | 17 | 2.95% | 80 | 17 | 5.35% |
| chr3 | 58 | 11 | 5.57% | 60 | 11 | 6.01% | 59 | 11 | 7.04% |
| chr4 | 65 | 19 | 5.11% | 71 | 19 | 6.85% | 67 | 19 | 5.90% |
| chr5 | 52 | 9 | 3.38% | 58 | 9 | 4.53% | 56 | 9 | 4.66% |
| chr6 | 55 | 13 | 2.52% | 60 | 14 | 4.39% | 59 | 13 | 6.02% |
| chr7 | 57 | 12 | 6.58% | 61 | 12 | 6.78% | 59 | 12 | 7.47% |
| chr8 | 46 | 12 | 4.34% | 47 | 12 | 5.40% | 47 | 12 | 4.84% |
| chr9 | 37 | 11 | 7.12% | 45 | 11 | 8.80% | 38 | 11 | 7.38% |
| chr10 | 54 | 15 | 5.15% | 56 | 15 | 5.73% | 56 | 16 | 7.04% |
| chr11 | 61 | 15 | 6.50% | 68 | 15 | 8.94% | 64 | 15 | 7.92% |
| chr12 | 45 | 15 | 3.71% | 47 | 15 | 4.37% | 47 | 15 | 5.64% |
| chr13 | 29 | 6 | 6.25% | 31 | 6 | 6.52% | 31 | 6 | 6.53% |
| chr14 | 28 | 5 | 5.97% | 30 | 5 | 6.45% | 27 | 5 | 6.88% |
| chr15 | 34 | 13 | 1.39% | 35 | 13 | 2.10% | 35 | 13 | 2.11% |
| chr16 | 34 | 10 | 2.84% | 34 | 10 | 2.84% | 35 | 10 | 4.09% |
| chr17 | 38 | 10 | 2.20% | 38 | 10 | 2.57% | 39 | 10 | 6.01% |
| chr18 | 31 | 5 | 5.86% | 33 | 5 | 5.87% | 31 | 5 | 6.63% |
| chr19 | 21 | 6 | 0.28% | 22 | 7 | 0.32% | 21 | 6 | 0.29% |
| chr20 | 17 | 4 | 4.65% | 18 | 4 | 6.38% | 17 | 4 | 10.14% |
| chr21 | 17 | 7 | 5.41% | 18 | 7 | 5.41% | 19 | 7 | 11.63% |
| chr22 | 13 | 4 | 2.10% | 13 | 4 | 3.64% | 13 | 4 | 3.64% |

**Table S18** N50 and Phased Block N50 of HapCUT2, MethPhaser, and HapBridge on ONT R9 60× Data

| R9 60 | HapCUT2 | MethPhaser | HapBridge | HapCUT2 | MethPhaser | HapBridge |
|---|---|---|---|---|---|---|
| | N50 | N50 | N50 | phase blockN50 | phase blockN50 | phase blockN50 |
| chr1 | 1872656 | 2058602 | 2676656 | 1671783 | 1823748 | 2340506 |
| chr2 | 1841365 | 2126285 | 2232599 | 1659141 | 1799794 | 2088423 |
| chr3 | 2499005 | 3018632 | 3187963 | 2209051 | 2675887 | 2939032 |
| chr4 | 2522287 | 2998771 | 3511966 | 2172950 | 2237396 | 3029796 |
| chr5 | 3348909 | 4751916 | 3845781 | 2368969 | 3562689 | 3562689 |
| chr6 | 1941674 | 2083306 | 2641752 | 1934688 | 2056055 | 2573352 |
| chr7 | 2434611 | 3064766 | 3053059 | 1912850 | 2398386 | 2304934 |
| chr8 | 2083284 | 2384385 | 3078034 | 1868720 | 2059575 | 2806414 |
| chr9 | 1823373 | 2141723 | 2141723 | 1687782 | 2123970 | 2123970 |
| chr10 | 1917053 | 2102143 | 2919419 | 1689854 | 1904929 | 2572148 |
| chr11 | 1986403 | 2919429 | 2810294 | 1969749 | 2689095 | 2792206 |
| chr12 | 2270793 | 3283800 | 3171502 | 1838466 | 2539921 | 2896786 |
| chr13 | 2009136 | 2690660 | 2398175 | 1760798 | 2188854 | 2103906 |
| chr14 | 2235100 | 2396319 | 3001288 | 2233504 | 2388648 | 2980414 |
| chr15 | 1520912 | 1738515 | 1696490 | 1309653 | 1663134 | 1620610 |
| chr16 | 1717691 | 2716973 | 3497585 | 1675690 | 2451452 | 2966943 |
| chr17 | 1745764 | 2087829 | 2986637 | 1652784 | 2081194 | 2985152 |
| chr18 | 1433525 | 1598683 | 2018016 | 1326423 | 1433525 | 1731210 |
| chr19 | 2300226 | 2626113 | 6089435 | 1940523 | 1940523 | 4045588 |
| chr20 | 2995359 | 3171943 | 3715991 | 2454551 | 2993226 | 3170063 |
| chr21 | 1102027 | 1102027 | 1247607 | 1036190 | 1036190 | 1232903 |
| chr22 | 1179053 | 1246285 | 2736610 | 1139620 | 1206852 | 2736266 |

**Table S19** N50 and Phased Block N50 of HapCUT2, MethPhaser, and HapBridge on R10 ONT 30× Data

| R10 30 | HapCUT2 N50 | MethPhaser N50 | HapBridge N50 | HapCUT2 phase blockN50 | MethPhaser phase blockN50 | HapBridge phase blockN50 |
|---|---|---|---|---|---|---|
| chr1 | 1655865 | 2480930 | 2169054 | 1620170 | 2375412 | 2030706 |
| chr2 | 1445907 | 1794190 | 2267211 | 1408643 | 1640637 | 2041931 |
| chr3 | 2055659 | 2446686 | 3167378 | 2046822 | 2401968 | 3161538 |
| chr4 | 2153300 | 2363332 | 2987866 | 1942018 | 2316814 | 2326850 |
| chr5 | 2850283 | 3491957 | 3296079 | 2560564 | 3252886 | 3070019 |
| chr6 | 2334507 | 3281061 | 2855076 | 1877405 | 2134697 | 2485556 |
| chr7 | 2206010 | 2456037 | 2464370 | 2205850 | 2412708 | 2427665 |
| chr8 | 1695451 | 2371899 | 2088736 | 1654096 | 2113302 | 2017396 |
| chr9 | 1792181 | 1792181 | 3099676 | 1581961 | 1615151 | 2710600 |
| chr10 | 1910549 | 2142552 | 2317726 | 1894013 | 2014888 | 2132437 |
| chr11 | 2011785 | 2835513 | 2672082 | 1933004 | 2629029 | 2593608 |
| chr12 | 2199437 | 2686781 | 2858401 | 2195439 | 2686691 | 2857762 |
| chr13 | 1668367 | 2130897 | 2604487 | 1619898 | 1808738 | 2098723 |
| chr14 | 2260777 | 2832565 | 2832565 | 2244257 | 2553085 | 2832559 |
| chr15 | 1311829 | 1462228 | 1738515 | 1096437 | 1311645 | 1571461 |
| chr16 | 2283046 | 3444735 | 2969232 | 2129082 | 2400305 | 2573680 |
| chr17 | 1593045 | 2342995 | 2608829 | 1577466 | 2342995 | 2583109 |
| chr18 | 1433525 | 1549234 | 1600203 | 1339823 | 1522999 | 1600197 |
| chr19 | 2575713 | 3850405 | 3727288 | 2384069 | 2891585 | 2384069 |
| chr20 | 2693221 | 2953670 | 3812480 | 2678355 | 2951537 | 3809167 |
| chr21 | 1110482 | 1110482 | 1619799 | 958970 | 1106026 | 1371748 |
| chr22 | 1233517 | 1307895 | 1643569 | 1232520 | 1307561 | 1447933 |

**Table S20** N50 and Phased Block N50 of HapCUT2, MethPhaser, and HapBridge on R10 ONT 60× Data

| R10 60 | HapCUT2 N50 | MethPhaser N50 | HapBridge N50 | HapCUT2 phase blockN50 | MethPhaser phase blockN50 | HapBridge phase blockN50 |
|---|---|---|---|---|---|---|
| chr1 | 2116144 | 2455501 | 3381001 | 2066966 | 2382548 | 3376334 |
| chr2 | 2043970 | 2839270 | 3601434 | 1950279 | 2655322 | 3595568 |
| chr3 | 2856802 | 3885940 | 3943442 | 2814196 | 3875862 | 3943265 |
| chr4 | 2610594 | 3606288 | 4391525 | 2588478 | 3269216 | 3440983 |
| chr5 | 2850283 | 3274464 | 5366354 | 2589423 | 2850041 | 4154713 |
| chr6 | 2651811 | 3599504 | 4190885 | 2647474 | 3577859 | 3944367 |
| chr7 | 2658749 | 3106590 | 3664818 | 2285546 | 2854792 | 3483636 |
| chr8 | 1991431 | 2597610 | 2606645 | 1953362 | 2565295 | 2576723 |
| chr9 | 2294843 | 2764822 | 5312851 | 2293287 | 2608952 | 4090791 |
| chr10 | 2284991 | 4414254 | 4401150 | 2133216 | 4202345 | 4202345 |
| chr11 | 2835513 | 3006480 | 3821992 | 2629029 | 2637646 | 3816631 |
| chr12 | 2813127 | 2948509 | 3292491 | 2689502 | 2879109 | 3149626 |
| chr13 | 1996145 | 3776105 | 3732521 | 1995859 | 3775840 | 3722180 |
| chr14 | 2969733 | 3086368 | 3677230 | 2962534 | 3085246 | 3643547 |
| chr15 | 1865322 | 2470880 | 2519650 | 1601873 | 2263802 | 1913631 |
| chr16 | 2969232 | 3452085 | 4356514 | 2701361 | 3044557 | 4307365 |
| chr17 | 2390810 | 2504187 | 3091804 | 2390281 | 2497122 | 3091799 |
| chr18 | 1653658 | 1777912 | 2193183 | 1652485 | 1770965 | 2045313 |
| chr19 | 2940589 | 4640468 | 6991484 | 2823018 | 2940589 | 3164014 |
| chr20 | 2973773 | 3361247 | 7169788 | 2962645 | 3359603 | 7167689 |
| chr21 | 1372483 | 1551125 | 2281805 | 1371933 | 1371933 | 1721103 |
| chr22 | 1506047 | 1506047 | 2739345 | 1504702 | 1506047 | 2739243 |

**Table S21** N50 and Phased Block N50 of HapCUT2, MethPhaser, and HapBridge on HiFi 30× Data

| HiFi 30 | HapCUT2 | | | MethPhaser | | | HapBridge | | |
|---|---|---|---|---|---|---|---|---|---|
| | SE | FE | HE | SE | FE | HE | SE | FE | HE |
| chr1 | 72 | 26 | 0.47% | 93 | 26 | 0.85% | 77 | 26 | 1.52% |
| chr2 | 57 | 18 | 0.54% | 78 | 18 | 1.23% | 64 | 18 | 0.97% |
| chr3 | 42 | 15 | 0.60% | 59 | 15 | 1.13% | 48 | 15 | 1.21% |
| chr4 | 62 | 24 | 0.61% | 87 | 24 | 2.35% | 63 | 24 | 0.78% |
| chr5 | 43 | 13 | 0.87% | 57 | 13 | 1.20% | 47 | 13 | 2.21% |
| chr6 | 49 | 17 | 0.35% | 73 | 18 | 0.81% | 50 | 17 | 1.21% |
| chr7 | 53 | 14 | 0.91% | 79 | 16 | 3.87% | 55 | 14 | 0.95% |
| chr8 | 41 | 14 | 0.49% | 52 | 14 | 0.72% | 47 | 14 | 2.71% |
| chr9 | 34 | 12 | 1.24% | 48 | 12 | 1.38% | 35 | 12 | 1.25% |
| chr10 | 66 | 21 | 2.92% | 72 | 21 | 3.25% | 69 | 21 | 4.88% |
| chr11 | 48 | 14 | 0.44% | 60 | 14 | 0.88% | 50 | 14 | 0.47% |
| chr12 | 48 | 16 | 2.20% | 56 | 17 | 3.19% | 49 | 16 | 2.74% |
| chr13 | 25 | 7 | 0.68% | 33 | 7 | 1.10% | 27 | 7 | 0.72% |
| chr14 | 15 | 5 | 0.55% | 22 | 5 | 1.06% | 16 | 5 | 0.56% |
| chr15 | 40 | 14 | 2.05% | 45 | 14 | 2.29% | 41 | 14 | 2.05% |
| chr16 | 30 | 9 | 0.82% | 40 | 9 | 1.96% | 35 | 9 | 1.28% |
| chr17 | 34 | 13 | 1.42% | 43 | 13 | 4.13% | 38 | 13 | 1.61% |
| chr18 | 25 | 7 | 2.02% | 31 | 7 | 3.20% | 27 | 7 | 2.03% |
| chr19 | 22 | 7 | 1.29% | 30 | 7 | 3.18% | 22 | 7 | 1.30% |
| chr20 | 12 | 4 | 0.87% | 17 | 4 | 4.76% | 13 | 4 | 1.54% |
| chr21 | 28 | 9 | 4.94% | 30 | 9 | 5.14% | 29 | 9 | 5.83% |
| chr22 | 10 | 4 | 0.16% | 17 | 4 | 2.26% | 10 | 4 | 0.16% |

**Table S22** N50 and Phased Block N50 of HapCUT2, MethPhaser, and HapBridge on HiFi 60× Data

| HiFi 60 | HapCUT2 | MethPhaser | HapBridge | HapCUT2 | MethPhaser | HapBridge |
|---|---|---|---|---|---|---|
| | N50 | N50 | N50 | phase blockN50 | phase blockN50 | phase blockN50 |
| chr1 | 518880 | 586238 | 599362 | 486643 | 552859 | 576045 |
| chr2 | 503156 | 582462 | 623037 | 493353 | 553744 | 594238 |
| chr3 | 665573 | 740794 | 773254 | 640092 | 671759 | 693098 |
| chr4 | 587940 | 695530 | 629810 | 577956 | 693567 | 615731 |
| chr5 | 678829 | 832089 | 799464 | 675486 | 773301 | 784564 |
| chr6 | 631466 | 695637 | 695637 | 571292 | 658111 | 641373 |
| chr7 | 630811 | 769029 | 784575 | 569521 | 710570 | 735935 |
| chr8 | 551107 | 693858 | 662365 | 546268 | 677106 | 657496 |
| chr9 | 461593 | 500890 | 480413 | 449967 | 470826 | 470274 |
| chr10 | 681134 | 714926 | 801753 | 628700 | 679783 | 727088 |
| chr11 | 524536 | 657747 | 628304 | 508439 | 627780 | 570824 |
| chr12 | 696392 | 878338 | 840925 | 673846 | 850438 | 791729 |
| chr13 | 479732 | 546694 | 511614 | 453440 | 526756 | 495972 |
| chr14 | 477781 | 501352 | 552752 | 461985 | 485290 | 526105 |
| chr15 | 336990 | 389103 | 413188 | 313615 | 362868 | 388077 |
| chr16 | 582558 | 693199 | 713652 | 564384 | 676395 | 681165 |
| chr17 | 492213 | 604133 | 672135 | 419237 | 551175 | 609510 |
| chr18 | 532512 | 613055 | 592055 | 514530 | 547876 | 547876 |
| chr19 | 720877 | 808249 | 1236533 | 598725 | 776334 | 1046467 |
| chr20 | 494333 | 531171 | 701128 | 480936 | 521066 | 691406 |
| chr21 | 197790 | 214943 | 210727 | 179531 | 209938 | 194026 |
| chr22 | 333782 | 409092 | 635809 | 306340 | 379667 | 533259 |

**Table S23** N50 and Phased Block N50 of HapCUT2, MethPhaser, and HapBridge on ONT R9 80× Data

| R9 80 | HapCUT2 | MethPhaser | HapBridge | HapCUT2 | MethPhaser | HapBridge |
|---|---|---|---|---|---|---|
| | N50 | N50 | N50 | phase blockN50 | phase blockN50 | phase blockN50 |
| chr1 | 2009584 | 2473100 | 3545417 | 1936280 | 2312358 | 3501887 |
| chr2 | 1955321 | 2481845 | 3025536 | 1799773 | 2284438 | 2864179 |
| chr3 | 2184549 | 2856802 | 4526145 | 2153387 | 2497470 | 4132876 |
| chr4 | 2695517 | 3634398 | 4439290 | 2458291 | 3067772 | 3913418 |
| chr5 | 3885850 | 3852318 | 7527461 | 3631613 | 3631613 | 7264009 |
| chr6 | 2239054 | 3101955 | 5047528 | 2056055 | 3094115 | 4598430 |
| chr7 | 2434611 | 3106590 | 3994270 | 1934377 | 2475480 | 3483636 |
| chr8 | 2233441 | 2526334 | 3964100 | 2041104 | 2320512 | 3838937 |
| chr9 | 3032767 | 3491785 | 4003233 | 2820289 | 3196611 | 4001056 |
| chr10 | 2096380 | 2616348 | 5054319 | 1934150 | 2356648 | 4600747 |
| chr11 | 2269371 | 3006480 | 4144462 | 2268611 | 2662191 | 3909081 |
| chr12 | 2751363 | 2975875 | 4675283 | 2130011 | 2172648 | 4064616 |
| chr13 | 2010698 | 3139456 | 4336377 | 1786457 | 2174853 | 3386224 |
| chr14 | 2295596 | 2498456 | 4142183 | 2294333 | 2490782 | 3358956 |
| chr15 | 1584185 | 1997160 | 3090510 | 1523803 | 1738473 | 3076916 |
| chr16 | 2283046 | 2969232 | 4874389 | 1733119 | 2282685 | 4743340 |
| chr17 | 2447535 | 3044237 | 5386783 | 1520285 | 2521291 | 4542327 |
| chr18 | 1528264 | 1853423 | 2793501 | 1293101 | 1827932 | 2780089 |
| chr19 | 2843916 | 4015181 | 15343139 | 2368685 | 2823018 | 8674175 |
| chr20 | 2456535 | 4629969 | 7085734 | 2454551 | 4628645 | 7085671 |
| chr21 | 1387846 | 1408996 | 1679614 | 1377605 | 1401988 | 1679038 |
| chr22 | 1179053 | 1611451 | 4033373 | 1139620 | 1598459 | 4033063 |

**Table S24** N50 and Phased Block N50 of HapCUT2, MethPhaser, and HapBridge on ONT R9 30× Data

| R9 30 | HapCUT2 | MethPhaser | HapBridge | HapCUT2 | MethPhaser | HapBridge |
|---|---|---|---|---|---|---|
| | N50 | N50 | N50 | phase blockN50 | phase blockN50 | phase blockN50 |
| chr1 | 1591955 | 1705195 | 1591955 | 1449430 | 1543203 | 1672531 |
| chr2 | 1355690 | 1491235 | 1376186 | 1308067 | 1409904 | 1672336 |
| chr3 | 1688343 | 1931849 | 1688343 | 1571185 | 1638659 | 1940230 |
| chr4 | 1698059 | 2234230 | 1724277 | 1498687 | 1883650 | 1714004 |
| chr5 | 2207887 | 2631862 | 2312302 | 2005979 | 2208049 | 2433253 |
| chr6 | 1484623 | 1648510 | 1484623 | 1355232 | 1470011 | 1919579 |
| chr7 | 1539335 | 1732189 | 1539335 | 1382327 | 1329790 | 1693163 |
| chr8 | 1619575 | 1840585 | 1619575 | 1478733 | 1685705 | 1667220 |
| chr9 | 1559365 | 1659555 | 1559365 | 1393278 | 1442410 | 1545021 |
| chr10 | 1440970 | 1546689 | 1477763 | 1264166 | 1426165 | 1871919 |
| chr11 | 1636379 | 1957504 | 1636379 | 1363267 | 1605488 | 1757645 |
| chr12 | 1563739 | 1809448 | 1563739 | 1465087 | 1559490 | 2018821 |
| chr13 | 1282848 | 1515496 | 1282848 | 1226756 | 1315832 | 1648838 |
| chr14 | 1378018 | 1575199 | 1378018 | 1314046 | 1384553 | 1554325 |
| chr15 | 1097623 | 1289902 | 1097623 | 858349 | 876198 | 1273559 |
| chr16 | 1737573 | 1968869 | 1960040 | 1732805 | 1737117 | 2712882 |
| chr17 | 1673994 | 1986099 | 1986099 | 1470774 | 1725023 | 2525063 |
| chr18 | 1471482 | 1768808 | 1471482 | 1201077 | 1365039 | 1433525 |
| chr19 | 1847732 | 2275421 | 1847732 | 1788610 | 2214464 | 2268695 |
| chr20 | 1463299 | 1570022 | 1463299 | 1327119 | 1461655 | 1922785 |
| chr21 | 636090 | 761784 | 636090 | 630911 | 711122 | 648461 |
| chr22 | 1179053 | 1332613 | 1179053 | 1003183 | 1025110 | 1623668 |

**Table S25** The N50 and Phased Block N50 of HapCUT2, MethPhaser, and HapBridge on NA12878 60× Data

| NA12878 60 | HapCUT2 | MethPhaser | HapBridge | HapCUT2 | MethPhaser | HapBridge |
|---|---|---|---|---|---|---|
| | N50 | N50 | N50 | phase blockN50 | phase blockN50 | phase blockN50 |
| chr1 | 660870 | 800910 | 759081 | 599442 | 709617 | 703435 |
| chr2 | 753292 | 913575 | 941362 | 664190 | 845205 | 843395 |
| chr3 | 843466 | 947036 | 948019 | 742204 | 881575 | 881575 |
| chr4 | 686828 | 855170 | 798520 | 628181 | 751551 | 707318 |
| chr5 | 840136 | 945007 | 957201 | 778887 | 871365 | 877656 |
| chr6 | 790185 | 919866 | 923903 | 743748 | 824266 | 836671 |
| chr7 | 838472 | 1067468 | 1115840 | 767918 | 893854 | 974588 |
| chr8 | 730103 | 937516 | 931872 | 700000 | 849217 | 849217 |
| chr9 | 660520 | 851206 | 864155 | 537300 | 677635 | 814769 |
| chr10 | 714149 | 932895 | 1077981 | 699709 | 861240 | 960113 |
| chr11 | 820104 | 1072090 | 1078110 | 720973 | 874196 | 885848 |
| chr12 | 722237 | 905535 | 1033066 | 698486 | 814970 | 925290 |
| chr13 | 578963 | 677338 | 677338 | 538794 | 591495 | 606867 |
| chr14 | 516665 | 709065 | 742123 | 467837 | 571581 | 693933 |
| chr15 | 365255 | 461568 | 541444 | 320173 | 415269 | 525089 |
| chr16 | 836707 | 1006883 | 1104107 | 746359 | 953783 | 944563 |
| chr17 | 561327 | 625109 | 899882 | 533072 | 608429 | 795163 |
| chr18 | 820004 | 853969 | 923010 | 789033 | 829664 | 882130 |
| chr19 | 1151568 | 1515589 | 2010553 | 1088907 | 1151566 | 1708017 |
| chr20 | 763232 | 925550 | 1004209 | 723314 | 843910 | 986565 |
| chr21 | 378833 | 492165 | 508419 | 343816 | 477404 | 492596 |
| chr22 | 654514 | 833035 | 833042 | 645423 | 793615 | 833035 |

**Table S26** The NG50 and Phased Block N50 of WhasHap, MethPhaser, and HapBridge

| | NG50 | | | Phased Block N50 | | |
|---|---|---|---|---|---|---|
| Sample | WhasHap | MethPhaser | HapBridge | WhasHap | MethPhaser | HapBridge |
| R9 80× | 2226283 | 2826074 | 4659112 | 1707881 | 2062157 | 3352926 |
| R9 60× | 1961289 | 2417775 | 2891363 | 1792404 | 2146139 | 2534741 |
| R9 30× | 1463934 | 1725904 | 2063936 | 1317983 | 1500602 | 1781226 |
| HiFi 60× | 534315 | 634024 | 659096 | 507520 | 567921 | 616791 |
| HiFi 30× | 412009 | 477709 | 493045 | 375455 | 423007 | 441005 |
| R10 60× | 4292226 | 4829306 | 6035782 | 3159013 | 3521557 | 4483625 |
| R10 30× | 1915265 | 2269755 | 3655007 | 1578033 | 1841095 | 2595596 |

**Table S27** The N50 and Phased Block N50 of HapCUT2, MethPhaser, and HapBridge on NA12878 60× Data

| NA12878 60 | WhasHap | MethPhaser | HapBridge | WhasHap | MethPhaser | HapBridge |
|---|---|---|---|---|---|---|
| | N50 | N50 | N50 | phase blockN50 | phase blockN50 | phase blockN50 |
| chr1 | 660137 | 753158 | 906840 | 599773 | 713788 | 781828 |
| chr2 | 745078 | 912511 | 1095728 | 663032 | 826673 | 995608 |
| chr3 | 843194 | 948019 | 1036252 | 736031 | 873800 | 917650 |
| chr4 | 686828 | 791250 | 855170 | 633936 | 714036 | 774602 |
| chr5 | 840334 | 1025555 | 1070975 | 775934 | 992448 | 992448 |
| chr6 | 790185 | 1053234 | 1104473 | 751229 | 938928 | 938928 |
| chr7 | 797825 | 1103745 | 1300295 | 767918 | 941902 | 1044175 |
| chr8 | 724593 | 927493 | 1056845 | 676176 | 764052 | 979861 |
| chr9 | 630469 | 697990 | 978729 | 579740 | 671937 | 855644 |
| chr10 | 714149 | 962395 | 1388338 | 699709 | 865273 | 1248313 |
| chr11 | 820104 | 923654 | 1123992 | 719241 | 813359 | 1055636 |
| chr12 | 722237 | 981914 | 1218053 | 696558 | 925290 | 1029012 |
| chr13 | 576531 | 705401 | 763334 | 566923 | 641244 | 708386 |
| chr14 | 514852 | 681866 | 912163 | 467837 | 626006 | 805960 |
| chr15 | 365255 | 461568 | 645024 | 353804 | 443266 | 634097 |
| chr16 | 836707 | 956152 | 1223686 | 790079 | 871984 | 1034059 |
| chr17 | 561327 | 651344 | 1033657 | 531605 | 618397 | 867172 |
| chr18 | 820004 | 949832 | 1108557 | 789033 | 915420 | 1031566 |
| chr19 | 1151568 | 1151568 | 2163688 | 1127963 | 1147076 | 2068116 |
| chr20 | 742842 | 1004209 | 1099063 | 723314 | 1002250 | 1037413 |
| chr21 | 378833 | 535561 | 609088 | 343816 | 489651 | 574878 |
| chr22 | 654514 | 926204 | 1027671 | 594077 | 825194 | 833035 |

**Table S28** The N50 and Phased Block N50 of WhasHap, MethPhaser, and HapBridge on ONT R9 30× Data

| R9 30 | WhasHap | MethPhaser | HapBridge | WhasHap | MethPhaser | HapBridge |
|---|---|---|---|---|---|---|
| | N50 | N50 | N50 | phase blockN50 | phase blockN50 | phase blockN50 |
| chr1 | 1490020 | 1705195 | 2055999 | 1398099 | 1463179 | 1731313 |
| chr2 | 1341977 | 1461643 | 1794190 | 1225309 | 1373130 | 1609260 |
| chr3 | 1656641 | 2024103 | 2367735 | 1571185 | 1669492 | 2032921 |
| chr4 | 1621085 | 1885609 | 1889012 | 1399583 | 1613591 | 1633120 |
| chr5 | 2149487 | 2312302 | 2942202 | 2064585 | 2149070 | 2675116 |
| chr6 | 1484623 | 1749597 | 2083915 | 1304429 | 1478017 | 1670866 |
| chr7 | 1490967 | 1693163 | 1774456 | 1322131 | 1477343 | 1544788 |
| chr8 | 1619575 | 1732247 | 1840585 | 1426255 | 1478733 | 1621643 |
| chr9 | 1480623 | 1824459 | 1824459 | 1427245 | 1480583 | 1566662 |
| chr10 | 1429208 | 1563072 | 2162589 | 1252872 | 1450953 | 1773223 |
| chr11 | 1392138 | 1913514 | 2036728 | 1334245 | 1528180 | 1757645 |
| chr12 | 1534604 | 1730913 | 2344520 | 1458465 | 1559490 | 1969407 |
| chr13 | 1282848 | 1379051 | 1778321 | 1126001 | 1226756 | 1522041 |
| chr14 | 1384188 | 1434706 | 1549023 | 1129630 | 1129630 | 1376936 |
| chr15 | 907915 | 1396171 | 1444646 | 858932 | 1273559 | 1302998 |
| chr16 | 1737573 | 2251819 | 3049232 | 1675690 | 2017356 | 2813828 |
| chr17 | 1673994 | 2460749 | 3127364 | 1417896 | 1797725 | 2465287 |
| chr18 | 1471482 | 1471482 | 2000775 | 1237026 | 1365039 | 1696878 |
| chr19 | 1847732 | 2214464 | 2385474 | 1631089 | 2197675 | 2274615 |
| chr20 | 1394733 | 1722869 | 2019876 | 1132085 | 1393741 | 1822724 |
| chr21 | 636090 | 761784 | 933539 | 599702 | 648461 | 702044 |
| chr22 | 1179053 | 1280987 | 2001973 | 1003183 | 1241554 | 1623668 |

**Table S29** The N50 and Phased Block N50 of WhasHap, MethPhaser, and HapBridge on ONT R9 60× Data

| R9 60 | WhasHap | MethPhaser | HapBridge | WhasHap | MethPhaser | HapBridge |
|---|---|---|---|---|---|---|
| | N50 | N50 | N50 | phase blockN50 | phase blockN50 | phase blockN50 |
| chr1 | 1872656 | 2120197 | 2687425 | 1707507 | 2032724 | 2340506 |
| chr2 | 1800287 | 2081401 | 2106750 | 1574892 | 1887654 | 1950279 |
| chr3 | 2214639 | 2704771 | 3110070 | 2055656 | 2387645 | 2624209 |
| chr4 | 2490843 | 3085516 | 2906808 | 2117502 | 2660856 | 2604104 |
| chr5 | 3296079 | 3514158 | 3845781 | 2881859 | 3079294 | 3348707 |
| chr6 | 1941674 | 2426188 | 2855287 | 1934688 | 2251543 | 2617516 |
| chr7 | 2017734 | 2840236 | 2927566 | 1858278 | 2433141 | 2433141 |
| chr8 | 2083284 | 2233441 | 3078034 | 1868720 | 2041104 | 2806414 |
| chr9 | 1823373 | 2141723 | 2141723 | 1687782 | 1998428 | 1998428 |
| chr10 | 1916769 | 2102143 | 2919419 | 1689854 | 1919850 | 2572148 |
| chr11 | 1990578 | 2545335 | 2810294 | 1969749 | 2300073 | 2802208 |
| chr12 | 2004366 | 2922123 | 3352749 | 1947586 | 2202408 | 3149626 |
| chr13 | 2009136 | 2189140 | 2398175 | 1575177 | 1760798 | 2103906 |
| chr14 | 1939204 | 2315377 | 3001288 | 1935147 | 2177327 | 2388648 |
| chr15 | 1520912 | 1677240 | 1696490 | 1309653 | 1539312 | 1539312 |
| chr16 | 1717691 | 2439910 | 3116873 | 1708705 | 2430868 | 2966943 |
| chr17 | 1660993 | 2504187 | 2909293 | 1603634 | 2183976 | 2897619 |
| chr18 | 1433525 | 1768808 | 2018016 | 1293101 | 1588462 | 1753650 |
| chr19 | 2184326 | 3530259 | 6089435 | 2083051 | 2590712 | 4493452 |
| chr20 | 2995359 | 3317456 | 3715991 | 2454551 | 3017456 | 3199779 |
| chr21 | 1055890 | 1387846 | 1247607 | 1036190 | 1387838 | 1232903 |
| chr22 | 1179053 | 1343595 | 2674928 | 1139620 | 1343595 | 1940809 |

**Table S30** The N50 and Phased Block N50 of WhasHap, MethPhaser, and HapBridge on HiFi 60× Data

| HiFi 60 | WhasHap | MethPhaser | HapBridge | WhasHap | MethPhaser | HapBridge |
|---|---|---|---|---|---|---|
| | N50 | N50 | N50 | phase blockN50 | phase blockN50 | phase blockN50 |
| chr1 | 516814 | 586238 | 587110 | 489507 | 552859 | 576045 |
| chr2 | 500190 | 582462 | 623037 | 490014 | 524301 | 580992 |
| chr3 | 665347 | 754786 | 708035 | 646533 | 691416 | 693098 |
| chr4 | 587940 | 719337 | 623297 | 577956 | 633949 | 612522 |
| chr5 | 678829 | 817270 | 790956 | 675486 | 781928 | 763784 |
| chr6 | 613488 | 723202 | 706871 | 562532 | 660358 | 657138 |
| chr7 | 630811 | 764672 | 784575 | 603296 | 678355 | 719491 |
| chr8 | 551107 | 693858 | 693858 | 543020 | 619200 | 649565 |
| chr9 | 462140 | 506207 | 500926 | 449967 | 461324 | 472388 |
| chr10 | 681134 | 725926 | 805042 | 615368 | 668021 | 751354 |
| chr11 | 524536 | 670608 | 646709 | 508439 | 529073 | 599164 |
| chr12 | 673867 | 891171 | 796640 | 673846 | 840202 | 791684 |
| chr13 | 479732 | 582103 | 511614 | 453440 | 493742 | 495972 |
| chr14 | 477781 | 606403 | 555467 | 461985 | 526105 | 551032 |
| chr15 | 336990 | 389103 | 412755 | 314841 | 339793 | 388077 |
| chr16 | 595467 | 713652 | 705365 | 575278 | 649081 | 676741 |
| chr17 | 485466 | 688271 | 673351 | 420158 | 564946 | 609442 |
| chr18 | 532512 | 579549 | 592055 | 514530 | 540434 | 547876 |
| chr19 | 720990 | 797861 | 1270996 | 613520 | 660152 | 1138479 |
| chr20 | 509993 | 562449 | 700342 | 482748 | 522297 | 623918 |
| chr21 | 197790 | 213448 | 203070 | 186655 | 194026 | 194026 |
| chr22 | 332016 | 379952 | 608058 | 306340 | 362704 | 476624 |

**Table S31** The N50 and Phased Block N50 of WhasHap, MethPhaser, and HapBridge on HiFi 30× Data

| HiFi 30 | WhasHap | MethPhaser | HapBridge | WhasHap | MethPhaser | HapBridge |
|---|---|---|---|---|---|---|
| | N50 | N50 | N50 | phase blockN50 | phase blockN50 | phase blockN50 |
| chr1 | 431736 | 512481 | 533317 | 375151 | 447786 | 470040 |
| chr2 | 412063 | 469126 | 476114 | 380183 | 424972 | 439635 |
| chr3 | 490278 | 587365 | 542210 | 477632 | 507785 | 492324 |
| chr4 | 466168 | 555828 | 553072 | 440078 | 478073 | 499109 |
| chr5 | 547863 | 605079 | 649805 | 485605 | 552514 | 557633 |
| chr6 | 466387 | 560038 | 528500 | 449653 | 502921 | 497779 |
| chr7 | 490720 | 548044 | 548044 | 442446 | 477603 | 505022 |
| chr8 | 479701 | 519318 | 551107 | 433655 | 471172 | 497196 |
| chr9 | 346848 | 411794 | 447829 | 317373 | 359332 | 367533 |
| chr10 | 547233 | 615646 | 644649 | 489066 | 536379 | 580063 |
| chr11 | 438961 | 472509 | 520574 | 384626 | 410825 | 477411 |
| chr12 | 496989 | 604790 | 551960 | 460647 | 519661 | 511701 |
| chr13 | 375763 | 469663 | 427817 | 345188 | 428121 | 368349 |
| chr14 | 348296 | 378582 | 405740 | 336722 | 360357 | 378060 |
| chr15 | 256194 | 311301 | 345499 | 223205 | 255843 | 298600 |
| chr16 | 401991 | 468267 | 511410 | 363774 | 410860 | 443689 |
| chr17 | 321288 | 381371 | 409445 | 282719 | 324095 | 353012 |
| chr18 | 419075 | 486690 | 502969 | 368936 | 448242 | 456877 |
| chr19 | 551705 | 600797 | 674359 | 518531 | 535490 | 582282 |
| chr20 | 362251 | 449255 | 449255 | 356000 | 407677 | 409665 |
| chr21 | 172652 | 196342 | 190036 | 141962 | 170176 | 167267 |
| chr22 | 240050 | 305317 | 383300 | 186873 | 276281 | 348883 |

**Table S32** The N50 and Phased Block N50 of WhasHap, MethPhaser, and HapBridge on R9 80× Data

| R9 80 | WhasHap | MethPhaser | HapBridge | WhasHap | MethPhaser | HapBridge |
|---|---|---|---|---|---|---|
| | N50 | N50 | N50 | phase blockN50 | phase blockN50 | phase blockN50 |
| chr1 | 2009584 | 3380223 | 3554281 | 1727295 | 1963313 | 2638391 |
| chr2 | 1914364 | 2502032 | 2947035 | 1565820 | 1913691 | 2502032 |
| chr3 | 2184549 | 2662951 | 5163397 | 2032921 | 2430639 | 3741756 |
| chr4 | 2783883 | 3691412 | 4043140 | 1893260 | 2291384 | 2772395 |
| chr5 | 3885850 | 4844283 | 7527461 | 2311526 | 2906611 | 5690888 |
| chr6 | 2102083 | 2487807 | 5047528 | 1859297 | 2330495 | 3577859 |
| chr7 | 2434611 | 2760491 | 4537518 | 1742748 | 2249672 | 3133136 |
| chr8 | 2134429 | 2842318 | 4098824 | 1896056 | 2237981 | 2836053 |
| chr9 | 2766969 | 3099588 | 3491791 | 1823373 | 2037238 | 3155431 |
| chr10 | 2096380 | 2919419 | 4601475 | 1773223 | 2121234 | 3478567 |
| chr11 | 2011785 | 2345004 | 3909371 | 1831877 | 1943627 | 3215394 |
| chr12 | 2443336 | 2912713 | 4064937 | 1947586 | 2557194 | 3791736 |
| chr13 | 2010698 | 2387060 | 4548954 | 1250040 | 1400315 | 2653868 |
| chr14 | 2295596 | 2945853 | 4142183 | 2085030 | 2490782 | 3053293 |
| chr15 | 1584185 | 1696490 | 3128557 | 1302998 | 1421894 | 2000618 |
| chr16 | 2283046 | 4182510 | 5151091 | 1675690 | 2563333 | 3359258 |
| chr17 | 2447535 | 2909293 | 4803213 | 1394014 | 1445879 | 3304931 |
| chr18 | 1528264 | 1732293 | 3993575 | 1201256 | 1293101 | 3074630 |
| chr19 | 2843916 | 4015181 | 10343139 | 2110056 | 2509017 | 5697431 |
| chr20 | 2456535 | 2857426 | 7085734 | 2004942 | 2568671 | 5430080 |
| chr21 | 1387846 | 1387846 | 2283892 | 1092940 | 1092940 | 2227610 |
| chr22 | 1372792 | 1611451 | 4033373 | 1051446 | 1598459 | 2429020 |

**Table S33** The N50 and Phased Block N50 of WhasHap, MethPhaser, and HapBridge on R10 60× Data

| R10 60 | WhasHap | MethPhaser | HapBridge | WhasHap | MethPhaser | HapBridge |
|---|---|---|---|---|---|---|
| | N50 | N50 | N50 | phase blockN50 | phase blockN50 | phase blockN50 |
| chr1 | 3673746 | 4393814 | 6041762 | 3150362 | 3671436 | 5238507 |
| chr2 | 3018224 | 3347067 | 4889878 | 2501585 | 2999098 | 4289085 |
| chr3 | 4791247 | 5260424 | 5746787 | 4285277 | 4317118 | 4983082 |
| chr4 | 5128765 | 5544449 | 7136082 | 4385472 | 4438766 | 5163245 |
| chr5 | 5185610 | 7055643 | 6492812 | 3369084 | 3452622 | 5050609 |
| chr6 | 3577942 | 4496835 | 4855936 | 2927541 | 3436451 | 4182951 |
| chr7 | 3664818 | 4361014 | 5771995 | 2836739 | 3209216 | 3749149 |
| chr8 | 4516292 | 5184956 | 7820138 | 2454443 | 3468215 | 3804483 |
| chr9 | 5963558 | 5963558 | 6198656 | 4059354 | 4059354 | 6196241 |
| chr10 | 3238450 | 4601839 | 5078853 | 2473789 | 3350235 | 4317232 |
| chr11 | 4094429 | 5560519 | 6395031 | 2798548 | 3008208 | 3109879 |
| chr12 | 4064937 | 4399343 | 4741388 | 3189075 | 3605027 | 3959867 |
| chr13 | 3047541 | 3263377 | 3908568 | 3016443 | 3263377 | 3732521 |
| chr14 | 4481902 | 6250665 | 6428220 | 3201404 | 5678123 | 5445661 |
| chr15 | 3114796 | 3155371 | 3412487 | 2161679 | 2348469 | 3155189 |
| chr16 | 7310187 | 7302270 | 10729317 | 3610408 | 3610408 | 6218035 |
| chr17 | 4354879 | 4376875 | 4376875 | 3062950 | 3261609 | 3261609 |
| chr18 | 4374715 | 4907368 | 5627792 | 1731586 | 2542833 | 2347424 |
| chr19 | 8406835 | 8406835 | 12715806 | 6935994 | 6935994 | 9022057 |
| chr20 | 4460006 | 4459867 | 7169788 | 3848083 | 3319234 | 6629630 |
| chr21 | 1785794 | 1785794 | 3208634 | 1598671 | 1598671 | 2486614 |
| chr22 | 2174319 | 2166851 | 4040401 | 1899800 | 1899800 | 2296699 |

**Table S34** The N50 and Phased Block N50 of WhasHap, MethPhaser, and HapBridge on R10 30× Data

| R10 30 | WhasHap | MethPhaser | HapBridge | WhasHap | MethPhaser | HapBridge |
|---|---|---|---|---|---|---|
| | N50 | N50 | N50 | phase blockN50 | phase blockN50 | phase blockN50 |
| chr1 | 1655865 | 2127114 | 3609297 | 1469639 | 1707585 | 2653144 |
| chr2 | 1432810 | 1783743 | 2861992 | 1362223 | 1461925 | 2249677 |
| chr3 | 2055659 | 2430217 | 3885940 | 1871270 | 2055659 | 3475895 |
| chr4 | 2153300 | 2363332 | 3293955 | 1690250 | 1774763 | 2876216 |
| chr5 | 2687840 | 2896752 | 4005548 | 2031412 | 2727390 | 3190499 |
| chr6 | 2171435 | 2485556 | 3969649 | 1679582 | 2046671 | 2921038 |
| chr7 | 2206010 | 2542088 | 4092789 | 1695828 | 2114800 | 2635316 |
| chr8 | 1695451 | 2083488 | 3938606 | 1407853 | 1654096 | 2384189 |
| chr9 | 1792181 | 1879967 | 3181162 | 1454997 | 1557814 | 2159290 |
| chr10 | 1910549 | 2015268 | 3344307 | 1703812 | 1792238 | 2290587 |
| chr11 | 2011785 | 2835513 | 3450925 | 1626774 | 1865518 | 2652098 |
| chr12 | 2199437 | 2686781 | 4290782 | 1934380 | 2393236 | 3791736 |
| chr13 | 1668367 | 1987759 | 3134135 | 1619898 | 1808738 | 2713137 |
| chr14 | 2260777 | 2832565 | 3360676 | 2233504 | 2648756 | 2832559 |
| chr15 | 1311829 | 1355857 | 1865322 | 862608 | 1023252 | 1571461 |
| chr16 | 2283046 | 2440946 | 4910721 | 1680465 | 2028428 | 2676261 |
| chr17 | 1593045 | 2342995 | 2940847 | 1369024 | 1562559 | 2123507 |
| chr18 | 1433525 | 1572080 | 2135674 | 1232242 | 1408575 | 1677675 |
| chr19 | 2575713 | 3354956 | 5251691 | 1551448 | 2384069 | 2932985 |
| chr20 | 2693221 | 3269384 | 5493087 | 2568879 | 2568879 | 3511281 |
| chr21 | 1110482 | 1110482 | 2059912 | 686633 | 686633 | 1170504 |
| chr22 | 1233517 | 1537770 | 5333139 | 984014 | 1232520 | 2614066 |

Table S9 and Table S26 present the results of WhasHap, MethPhaser, HapBridge on different sequencing datasets. Table S18 compares the performance of three genome assembly tools under different sample types (R9, R10, HiFi) and sequencing depths (30×, 60×, 80×) using two key metrics: NG50 and Phased Block N50. HapBridge demonstrated a significant advantage in NG50, consistently outperforming WhasHap and MethPhaser across all sample types and sequencing depths. For instance, in the R10 60× sample, HapBridge achieved an NG50 of 6,035,782, which is 25% and 40% higher than the NG50 values of MethPhaser (4,829,306) and WhasHap (4,292,226), respectively. Consistent with the experimental results for HapCUT2, the N50 values of phasing algorithms increased progressively with higher sequencing depths. HapBridge also exhibited the best performance in Phased Block N50. For example, in the R9 80× sample, HapBridge achieved a Phased Block N50 of 3,352,926, significantly exceeding the values of MethPhaser (2,062,157) and WhasHap (1,707,881).

Table S34 presents a comparison of Switch Error and Flip Error metrics among the three tools. HapBridge demonstrated superior performance in Switch Error, particularly at sequencing depths exceeding 60×, with values significantly lower than those of the other two tools. For instance, in the R9 80× sample, HapBridge achieved a Switch Error of 658, compared to 717 for WhasHap and 758 for MethPhaser. The performance of the three tools was comparable in terms of Flip Error across all conditions.

**Table S35** The N50 values of MARGIN, HapBridge, and HapCUT2 across HiFi 60×, R9 30×, and R9 60× datasets.

| | HiFi 60× | R9 30× | R9 60× |
|---|---|---|---|
| MARGIN | 222986 | 1465398 | 1881717 |
| MARGIN-HapBridge | 259372 | 1749245 | 2524651 |
| HapCUT2 | 414153 | 1506983 | 2035463 |
| HapCUT2-HapBridge | 507697 | 2170238 | 2939026 |

Among the MARGIN-based phasing methods, as shown in Table, HapBridge still improves phasing performance; however, its overall results are limited by the relatively low baseline performance of MARGIN itself when compared to HapCUT2. This limitation is particularly evident in the HiFi dataset, where the N50 of phased blocks produced by MARGIN is only about half of that achieved by HapCUT2. MARGIN is a suite of tools that performs long-read data analysis based on Hidden Markov Models. As a result, its performance on HiFi data is suboptimal, and its typical usage scenario is within the PEPPER-Margin-DeepVariant variant calling pipeline, rather than being employed as a standalone SNP phasing tool.

**Table S36** Genome assembly performance comparison across different phasing strategies on R9, HiFi, and R10 datasets. The Block N50, Switch Error (%), and Hamming Error (%) for four methods: Hapdup-HapCUT2, Hapdup-HapCUT2 (HapBridge).

| DATA | Method | Block N50 | switch error(%) | hamming_error(%) |
|---|---|---|---|---|
| | | dual1\|dual2 | dual1\|dual2 | dual1\|dual2 |
| R9 80x | HapDup-HapCUT2 | 3615109\|3600776 | 0.159554\|0.156770 | 21.7065\|21.6443 |
| | HapDup-HapCUT2(HapBridge) | 5359421\|5602931 | 0.155656\|0.151975 | 19.1712\|19.1636 |
| HiFi 60x | HapDup-HapCUT2 | 1344264\|1349350 | 0.104925\|0.106300 | 29.2420\|28.9484 |
| | HapDup-HapCUT2(HapBridge) | 1515650\|1516986 | 0.097513\|0.095984 | 28.8561\|28.6145 |
| R10 30x | HapDup-HapCUT2 | 3823458\|3720602 | 0.067722\|0.060815 | 21.2823\|21.3359 |
| | HapDup-HapCUT2(HapBridge) | 5480848\|5498163 | 0.065179\|0.059371 | 18.2681\|19.0249 |

**Table S37** The comparison of the number of highly consistent methylation sites and haplotype-specific methylation sites on Chr1 in ONT R9, R10, and HiFi data.

| | num_haplotype-specific methylation | num_highly consistent methylation | num_overlap | Overlap rate |
|---|---|---|---|---|
| R9 | 225946 | 67699 | 58134 | 85.87% |
| R10 | 226325 | 57943 | 50624 | 87.37% |
| HiFi | 175863 | 65777 | 50880 | 77.35% |

Table S37 presents the overlap between highly consistent methylation sites and haplotype-specific methylation sites, which is intended to demonstrate the accuracy of consistent methylation sites in the absence of haplotype information. We assume that haplotype-specific methylation sites are fully accurate, and thus, the overlap can be interpreted as the accuracy. In the R9 data, the accuracy of methylation sites is relatively high, while the overlap rate in the HiFi data is the lowest, yet still reaches 77.35%. This indicates that, even without haplotype information, the accuracy of the highly consistent methylation sites we selected exceeds 70%.

**Table S38** HG001 (60× Coverage) phasing performance comparison: WhatsHap, HapCUT2, MethPhaser, and HapBridge evaluated by Switch Error, Flip Error, N50, and Phased Block N50

| Method | Switch Error | Flip Error | N50 | Phased Block N50 |
|---|---|---|---|---|
| HapCUT2 | 514 | 63 | 708071 | 663079 |
| HapCUT2-MethPhaser | 630 | 81 | 868392 | 800999 |
| HapCUT2-HapBridge | 606 | 102 | 1078256 | 964018 |
| WhatsHap | 646 | 102 | 713065 | 658080 |
| WhatsHap-MethPhaser | 818 | 111 | 874934 | 776882 |
| WhatsHap-HapBridge | 796 | 125 | 939693 | 851265 |

To further assess the generalizability of HapBridge, we evaluated its performance on the HG001 (NA12878) genome at 60× coverage and compared it with WhatsHap, HapCUT2, and MethPhaser. Consistent with results on HG002, HapBridge produced substantially higher N50 and phased-block N50 when paired with either WhatsHap or HapCUT2. As shown in Table 2, HapCUT2-HapBridge shows superior block continuity, with an N50 of 939,693 and a phased block N50 of 851,265, making it the best-performing method among the HapCUT2-based approaches. WhatsHap-HapBridge delivers the best overall performance, with substantial improvements in N50 (1,078,256) and phased block N50 (964,018) compared to WhatsHap and WhatsHap-MethPhaser. Although HapBridge introduces a modest increase in switch-error counts, this elevation is primarily attributable to flip errors rather than long-range switch errors, indicating that the improvements in haplotype contiguity are achieved without compromising global phasing accuracy. Overall, these findings demonstrate that HapBridge is robust across distinct SNP-based phasing tools and datasets, confirming that DNA methylation provides orthogonal and complementary linkage information that enhances long-read phasing independent of the underlying variant-calling approach.

**Table S38** Block N50, Switch Error (%), and Hamming Error (%) for Hapdup, HapBridge, and HapCUT2-Hapbridge on ONT R9 data (30×).

| **R9 30×** | **Block N50** | **switch error(%)** | **hamming_error(%)** |
|---|---|---|---|
| **Merqury** | **dual1\|dual2** | **dual1\|dual2** | **dual1\|dual2** |
| Hapdup | 2394535\|2311110 | 0.205391\|0.215488 | 22.52\|22.35 |
| HapBridge | 2457158\|2493891 | 0.214778\|0.217235 | 22.17\|22.50 |
| HapCUT2-Hapbridge | 2667695\|2536125 | 0.221817\|0.210876 | 22.12\|21.6 |

**Table S39** BlockN50, Switch Error (%), and Hamming Error (%) for Hapdup, HapBridge, and HapCUT2 Hapbridge on HiFi data (30×).

| **HiFi 30×** | **Block N50** | **switch error(%)** | **hamming_error(%)** |
|---|---|---|---|
| **Merqury** | **dual1\|dual2** | **dual1\|dual2** | **dual1\|dual2** |
| Hapdup | 1326150\|1337076 | 0.121412\|0.120968 | 22.3\|21.933 |
| HapBridge | 1427393\|1416290 | 0.121095\|0.115571 | 22.46\|21.90 |
| HapCUT2-Hapbridge | 1238138\|1226222 | 0.156214\|0.155196 | 26.90\|27.17 |

**Table S40** The performance of three different phasing algorithms during the Hapdup pipline.

| | R9 30× | HiFi 30× |
|---|---|---|
| Hapdup(MARGIN) | 1467191 | 603883 |
| HapBridge | 1749245 | 689336 |
| HapCUT2 | 1486747 | 433397 |
| HapBridge(HapCUT2) | 1876921 | 514128 |

Tables S38 and S39 present the results of HapBridge during the assembly process. On R9 30× data, the results show an improvement compared to Hapdup, whether based directly on Margin or on HapCUT2. However, on HiFi 30× data, particularly for experiments based on HapCUT2, the results deteriorate.

As shown in Table S40, the N50 metric improves on ONT R9 30× data. However, on HiFi data, the experimental results for both HapCUT2 and HapBridge (based on HapCUT2) are inferior to those of Margin. This further demonstrates that the phasing quality directly impacts the results of genome assembly.

**Table S41** HapBridge runtime

| Datasets | WhasHap | MethPhaser | HapCUT2 | HapBridge |
|---|---|---|---|---|
| R9 60× | 26,679 | 81,534 | 18,104 | 33,667 |
| R9 80× | 35,572 | 108,712 | 24,737 | 57,821 |
| R10 60× | 41,040 | 113,594 | 28,540 | 29,481 |
| HiFi 30× | 8,966 | 27,600 | 4,021 | 9,200 |
| R9 30× | 10,549 | 42,927 | 5,820 | 15,861 |
| R10 30× | 16,229 | 59,997 | 8,954 | 15,937 |

As shown in Table S41, WhasHap and HapCUT2, two SNP-based phasing tools, run the fastest, while MethPhaser runs the slowest. Moreover, the higher the data coverage, the longer the running time of the four algorithms. In short, HapBridge has an advantage over MethPhaser, but not over the other two phasing tools.

**Table S42** The N50 of MARGIN, HapBridge on HiFi 60×, R9 30× and R9 60× Data

| | HiFi 60× | R9 30× | R9 60× |
|---|---|---|---|
| MARGIN | 222986 | 1465398 | 1881717 |
| MARGIN-HapBridge | 259372 | 1749245 | 2524651 |
| MARGIN-MethPhaser | 241273 | 1596218 | 2318301 |

Table S42 presents a comparison of N50 values obtained using three different phasing methods—MARGIN, HapBridge, and MethPhaser—under three sequencing platforms and coverage conditions: HiFi 60×, R9 30×, and R9 60×. Among these methods, HapBridge demonstrates the best performance, consistently achieving higher N50 values across all datasets. For instance, in the R9 60× dataset, the N50 increased from 1,881,717 with MARGIN alone to 2,524,651 with MARGIN-HapBridge, representing an improvement of approximately 34.2%. While MethPhaser also contributed to improved phasing continuity, it performed slightly below HapBridge. In the R9 60× dataset, MARGIN-MethPhaser achieved an N50 of 2,318,301, which is higher than MARGIN alone (1,881,717) but lower than MARGIN-HapBridge (2,524,651). A similar trend was observed in the HiFi 60× dataset, where MethPhaser reached an N50 of 241,273, compared to 259,372 achieved by HapBridge.

**Table S43** Genome assembly performance comparison on the HG002 sample across different phasing methods using R9, HiFi, and R10 datasets. Phase Block N50, intra-block Switch Error (\%), and Hamming Error (\%) for three methods: HapDup, HapBridge, MethPhaser.

<table>
<tr><th>DATA</th><th>Method</th><th>Phase Block N50</th><th>intra-block Switch Error(%)</th><th>Hamming Error(%)</th></tr>
<tr><td></td><td></td><td>dual1|dual2</td><td>dual1|dual2</td><td>dual1|dual2</td></tr>
<tr><td rowspan="3">R9 80x</td><td>Hapdup-MARGIN</td><td>3714884|3673789</td><td>0.148158|0.161225</td><td>20.3932|20.2345</td></tr>
<tr><td>Hapdup-MARGIN(HapBridge)</td><td>4423667|4548083</td><td>0.156033|0.154542</td><td>18.6206|18.9672</td></tr>
<tr><td>Hapdup-MARGIN(MethPhaser)</td><td>4015109|3970776</td><td>0.161554|0.163770</td><td>21.2321|21.9553</td></tr>
<tr><td rowspan="3">HiFi 60x</td><td>Hapdup-MARGIN</td><td>650769|647135</td><td>0.109354|0.113797</td><td>25.6835|25.7419</td></tr>
<tr><td>Hapdup-MARGIN(HapBridge)</td><td>799211|796458</td><td>0.104239|0.106223</td><td>25.5809|25.5630</td></tr>
<tr><td>Hapdup-MARGIN(MethPhaser)</td><td>754214|750353</td><td>0.111531|0.113320</td><td>25.7022|25.7684</td></tr>
<tr><td rowspan="3">R10 30x</td><td>Hapdup-MARGIN</td><td>3796174|3747906</td><td>0.055526|0.061596</td><td>22.4862|22.3260</td></tr>
<tr><td>Hapdup-MARGIN(HapBridge)</td><td>4245293|4292473</td><td>0.058013|0.064249</td><td>20.0838|20.2777</td></tr>
<tr><td>Hapdup-MARGIN(MethPhaser)</td><td>4154890|4170213</td><td>0.057982|0.062190</td><td>21.1763|21.0057</td></tr>
</table>

### 2. Supplementary Figures

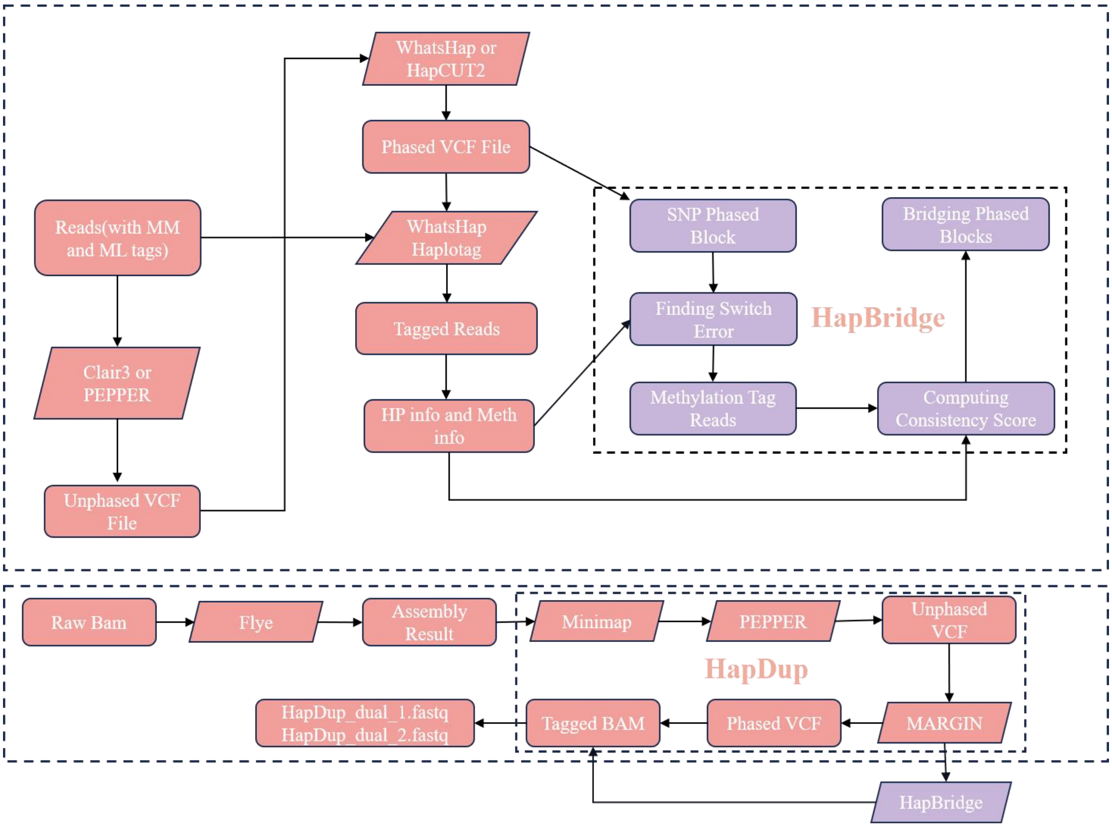


Figure S1 HapBridge's flowchart

Figure S1 illustrates the operational workflow of HapBridge. For an initial BAM file, the primary prerequisite is that the file contains methylation site information. Subsequently, it is necessary to call SNPs, for which tools such as Clair3 or PEPPER can be employed. These SNPs are then subjected to haplotype phasing using tools such as WhasHap, HapCUT2, or MARGIN. Following this, the **WhasHap haplotag** command is utilized to assign haplotype tags to the reads within the BAM file. Finally, the phased VCF file and the tagged BAM file serve as input files for HapBridge. The output of HapBridge consists of a VCF file and an optionally generated HapBridge-tagged BAM file.

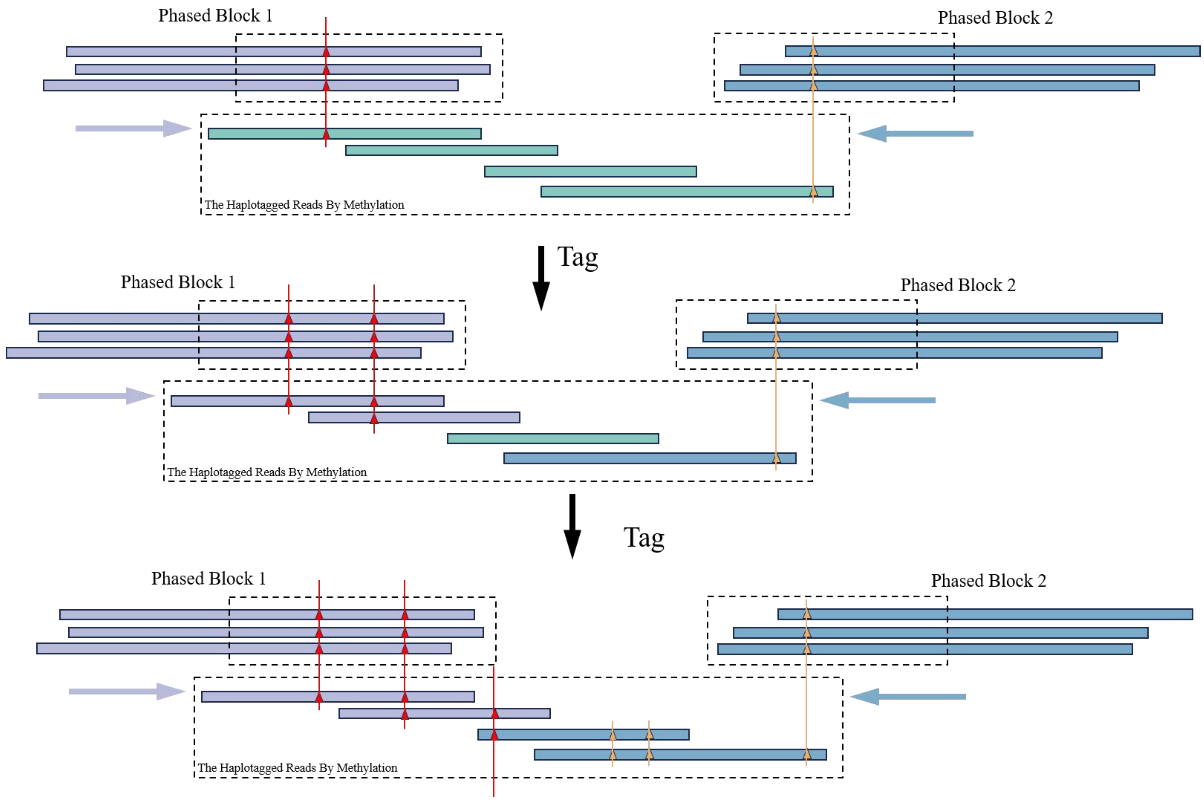


Figure S2 The process where reads from two blocks merge towards the center

Between the two phased blocks, we select two types of methylation sites. One type consists of methylation sites with high consistency scores that do not rely on haplotype information. The other type consists of haplotype-specific methylation sites identified during the extension process of the two blocks, where haplotype information is used; however, this haplotype is determined based on methylation and thus has lower accuracy compared to SNP-based haplotypes. To improve accuracy, we take the intersection of these two types of methylation sites as candidate sites for connecting the two blocks. In this way, by continuously performing haplotype determination on the reads between two Phased Blocks, candidate methylation sites for connecting the two Blocks are obtained. Phased Block 1 (blk1) and Phased Block 2 (blk2) will each obtain their respective candidate methylation sites. For each read passing through these methylation sites, we check whether the haplotypes of the two sets of candidate methylation sites are consistent across the read. If they are consistent, the two phased blocks can be directly connected; if not, the blocks are connected after being flipped. If no such candidate methylation sites exist, this indicates that the two blocks cannot be connected. The implementation method is as follows:

In two unlinked haplotype blocks, denoted as blk1 and blk2, HapBridge initially identifies a set of candidate methylation sites for bridging in prior steps. Subsequently, the algorithm iterates over all sequencing reads spanning the interval between these blocks. For each read, HapBridge independently determines the haplotype (HP) and phase set (PS) tags for blk0 and blk1 based on the identified methylation sites and evaluates the HP tag consistency. When the same read yields valid HP tag determinations for both blocks, the method compares whether these HP tags are identical. Finally, HapBridge outputs one of three possible outcomes: 1, indicating that the blocks can be directly bridged; −1, signifying that bridging is possible after a phase flip; or 0, denoting insufficient information to establish a connection.

### 3. Software Versions and Command Usage

**The version of WhasHap: 1.7**

Using WhasHap as an example, Clair3:latest was used for variant calling with the consideration of different read types, then WhasHap (WhasHap phase -o VCF--reference=REF input.vcf input.bam --ignore-read-groups) for SNV phasing, (WhasHap haplotag -o haplotagged.bam --reference reference.fasta phased.vcf.gz alignments.bam --ignore-read-groups) for read tag, (WhasHap compare --names truth,WhasHap --tsv-pairwise eval.tsv truth.chr1.vcf phased.chr1.vcf) for comparing variant files, and (WhasHap stats input.vcf) for computing N50.

**The version of HapCUT2: 1.3.4**

Github: https://github.com/vibansal/HapCUT2

./build/extractHAIRS [options] --bam reads.sorted.bam --VCF variants.vcf --out fragment_file

./build/HAPCUT2 --fragments fragment_file --VCF variants.vcf --output haplotype_output_file

**The version of MethPhaser:V0.0.3.**

Github: https://github.com/treangenlab/MethPhaser?tab=readme-ov-file

Usage: Strictly followed the instructions.

./meth_phaser_parallel -b test_data/HLA.R10.haplotagged.bam -r test_data/GCA_000001405.15_GRCh38_no_alt_analysis_set.chr6.fna -g test_data/LSK.filtered.gtf -vc test_data/HLA.R10.phased.vcf.gz -o test_data/work

./meth_phaser_post_processing -ib test_data/HLA.R10.haplotagged.bam -if test_data/work/ -ov test_data/output.vcf -ob test_data/output -vc test_data/HLA.R10.phased.vcf.gz

HapBridge: https://github.com/Humonex/HapBridge

**The version of Merqury :v1.3**

Github: [GitHub - marbl/merqury: k-mer based assembly evaluation]

Usage: merqury.sh <read-db.meryl> [<mat.meryl> <pat.meryl>] <asm1.fasta> [asm2.fasta] <out>

**The version of Hapdup: 0.12**

Github: https://github.com/KolmogorovLab/hapdup

**Hamming Error**: The computation of Hamming Error depends on the *bench.hapmers.count* file generated by **Merqury**, and the corresponding implementation code is shown below: HapBridge/Hamming Error at main · Humonex/HapBridge.

**Phased Block N50** : HapBridge/Phased Block N50 at main · Humonex/HapBridge

### 4. Confirmation of parameters in the paper

In the paper, we conducted the following experimental analysis on the three parameters involved: **Sup_count**, **Sup_rate**, and **Distance**.

***Distance***

To evaluate the impact of the **Distance** parameter on the identification of haplotype-specific methylation sites. Regarding the analysis of this value, we first extracted the number of methylation sites at different **Distance** values. As shown in the figure below, as the **Distance** increases, the number of filtered methylation sites decreases, but no clear pattern is observed.

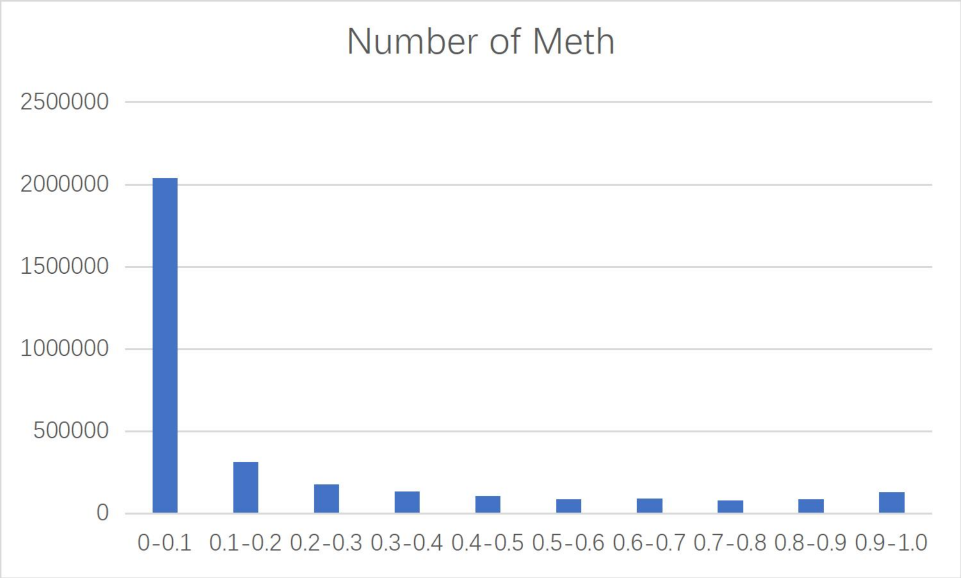


Figure S3 Number of Methylation Sites Retained at Different Distance Thresholds

For the HG002 R9 80× data, methylation sites and corresponding details were extracted at **Distance**= 0.1, 0.2, 0.3, 0.4, and 0.5 (see **Figure S3**).

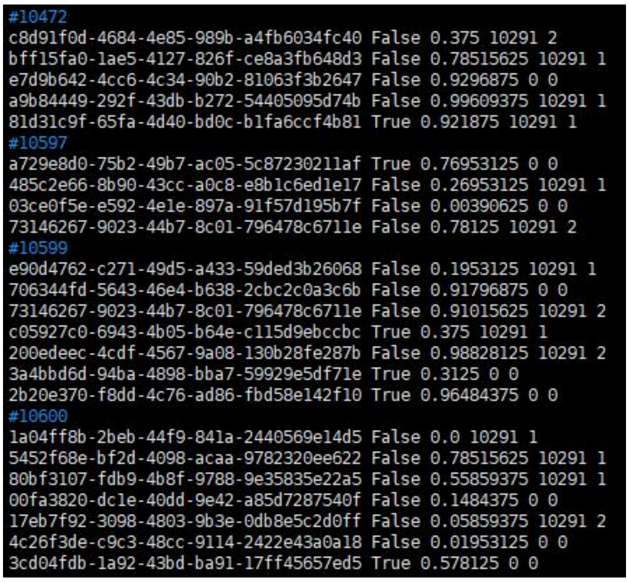

```
#10472
c8d91f0d-4684-4e85-989b-a4fb6034fc40 False 0.375 10291 2
bff15fa0-1ae5-4127-826f-ce8a3fb648d3 False 0.78515625 10291 1
e7d9b642-4cc6-4c34-90b2-81063f3b2647 False 0.9296875 0 0
a9b84449-292f-43db-b272-54405095d74b False 0.99609375 10291 1
81d31c9f-65fa-4d40-bd0c-b1fa6ccf4b81 True 0.921875 10291 1
#10597
a729e8d0-75b2-49b7-ac05-5c87230211af True 0.76953125 0 0
485c2e66-8b90-43cc-a0c8-e8b1c6ed1e17 False 0.26953125 10291 1
03ce0f5e-e592-4e1e-897a-91f57d195b7f False 0.00390625 0 0
73146267-9023-44b7-8c01-796478c6711e False 0.78125 10291 2
#10599
e90d4762-c271-49d5-a433-59ded3b26068 False 0.1953125 10291 1
706344fd-5643-46e4-b638-2cbc2c0a3c6b False 0.91796875 0 0
73146267-9023-44b7-8c01-796478c6711e False 0.91015625 10291 2
c05927c0-6943-4b05-b64e-c115d9ebccbc True 0.375 10291 1
200edeec-4cdf-4567-9a08-130b28fe287b False 0.98828125 10291 2
3a4bbd6d-94ba-4898-bba7-59929e5df71e True 0.3125 0 0
2b20e370-f8dd-4c76-ad86-fbd58e142f10 True 0.96484375 0 0
#10600
1a04ff8b-2beb-44f9-841a-2440569e14d5 False 0.0 10291 1
5452f68e-bf2d-4098-acaa-9782320ee622 False 0.78515625 10291 1
80bf3107-fdb9-4b8f-9788-9e35835e22a5 False 0.55859375 10291 1
00fa3820-dc1e-40dd-9e42-a85d7287540f False 0.1484375 0 0
17eb7f92-3098-4803-9b3e-0db8e5c2d0ff False 0.05859375 10291 2
4c26f3de-c9c3-48cc-9114-2422e43a0a18 False 0.01953125 0 0
3cd04fdb-1a92-43bd-ba91-17ff45657ed5 True 0.578125 0 0
```

Figure S4 Detailed Information on Filtered Methylation Sites

The variance of each methylation site was computed to assess the heterogeneity of methylation levels. Larger variances (with a maximum of 0.25 due to normalization) imply a bimodal distribution, consistent with haplotype-specific methylation. The average variance across all sites was then derived, where higher values suggest more pronounced haplotype-specific signals. The table below presents the variance of methylation sites selected at varying **Distance** values, reflecting their haplotype-specific differentiation potential.

**Table S44** The number of methylation sites screened under different Distance values and the mean of their variances.

| Distance | Number of Meth | Mean of the variance |
|---|---|---|
| 0.1 | 1217588 | 0.12852 |
| 0.2 | 902773 | 0.139457 |
| 0.3 | 725474 | 0.150294 |
| 0.4 | 591180 | 0.160801 |
| 0.5 | 484170 | 0.170172 |

The Table S44 indicates a clear positive correlation between the distance threshold and the mean variance value, though this relationship exhibits diminishing returns. Specifically:

1. While increasing the distance from 0.1 to 0.2 yields substantial gains in mean variance, the improvement becomes less pronounced beyond 0.3.
2. This plateau effect coincides with stabilization in the number of filtered methylation sites.
3. The inflection point at distance=0.3 represents an optimal trade-off between:

Therefore, we recommend distance=0.3 as the default threshold, providing 85% of the maximum variance benefit while retaining 70% more methylation sites compared to more stringent thresholds.

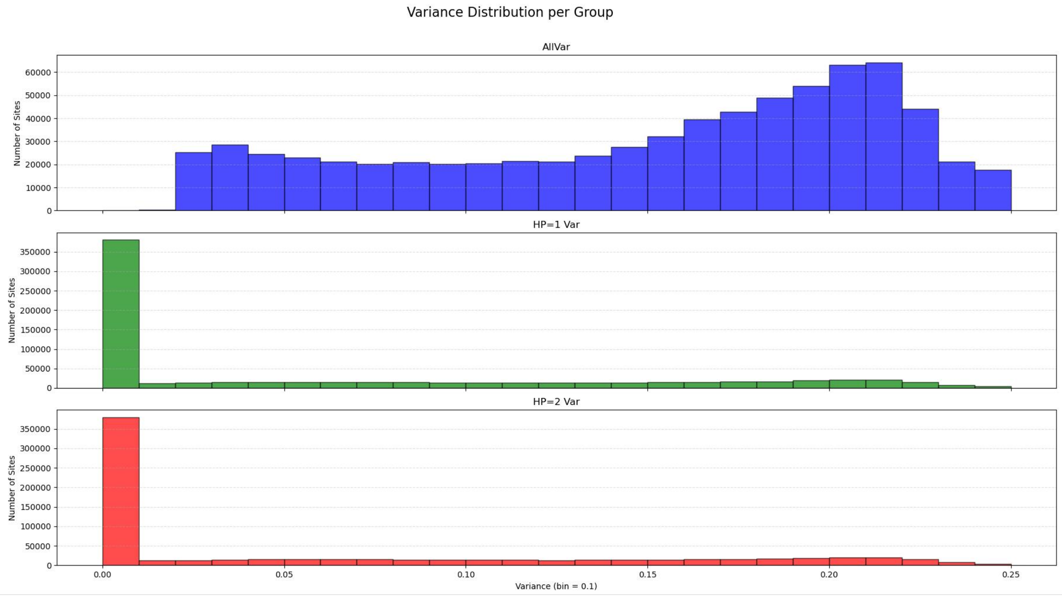


**Figure S5** Variance Distribution Plot of Methylation Sites (Distance = 0.3)

Figure S5 displays the variance distribution of the filtered methylation sites. The x-axis of this figure represents the variance values of methylation sites with a binning interval of 0.01, while the y-axis indicates the number of methylation sites, corresponding to the count of methylation sites within each variance range. The blue bars denote the variance of methylation values at these sites, the green bars represent the variance of methylation values labeled as HP=1, and the red bars indicate the variance of methylation values labeled as HP=2. As evident from the figure, the vast majority of the filtered methylation sites exhibit variances concentrated within the 0-0.01 range, indicating that these sites predominantly display either high or low methylation values, demonstrating distinct haplotype-specific characteristics.

***Sup_rate***

To evaluate the impact of the **Sup_rate** parameter on the identification of haplotype-specific methylation sites. Under the condition of **Distance = 0.3**, we analyzed the effect of varying **Sup_rate** values on the filtering of methylation sites. As illustrated in the figure below, the results demonstrate that when **Sup_rate ≥ 0.1** or **≥ 0.2**, the filtering efficiency of methylation sites remains moderate, with no significant difference in the number of retained sites between the **0.1–0.2** range. However, a noticeable reduction in methylation site counts becomes evident starting at **Sup_rate = 0.3**. Based on this observation, we conclude that **Sup_rate = 0.3** is an optimal parameter choice, as it effectively balances specificity and data retention while reflecting a clear inflection point in site exclusion.

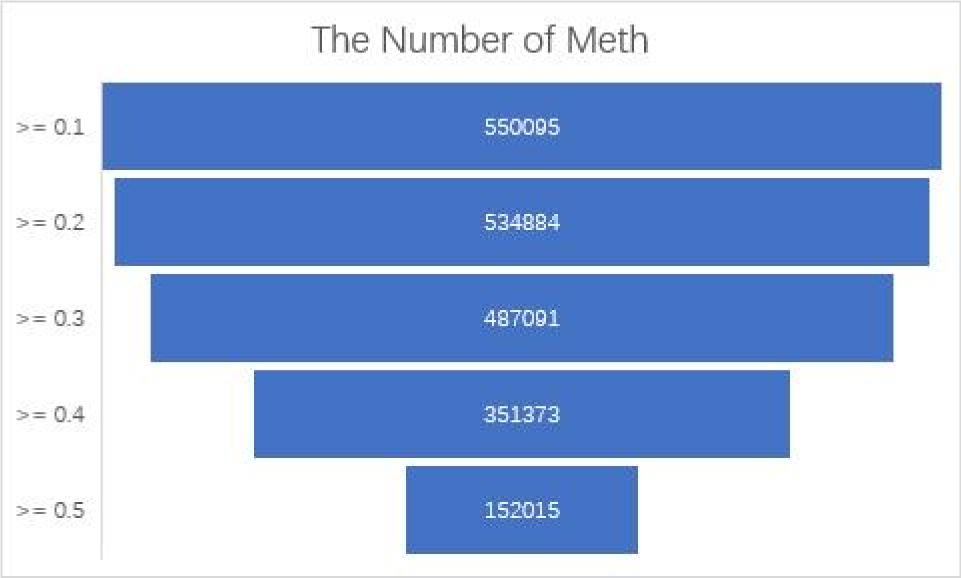


**Figure S6** Variance Distribution Plot of Methylation Sites

Meanwhile, we also examined the variance distribution of methylation sites under the conditions of **Distance=0.3** and **Sup_rate=0.3**, as illustrated in Figure S6. Comparative analysis with the initial figure reveals a pronounced enrichment of methylation sites with lower variances through the additional filtering step. Notably, the majority of methylation sites exhibit a clustered distribution in regions exceeding 0.15.

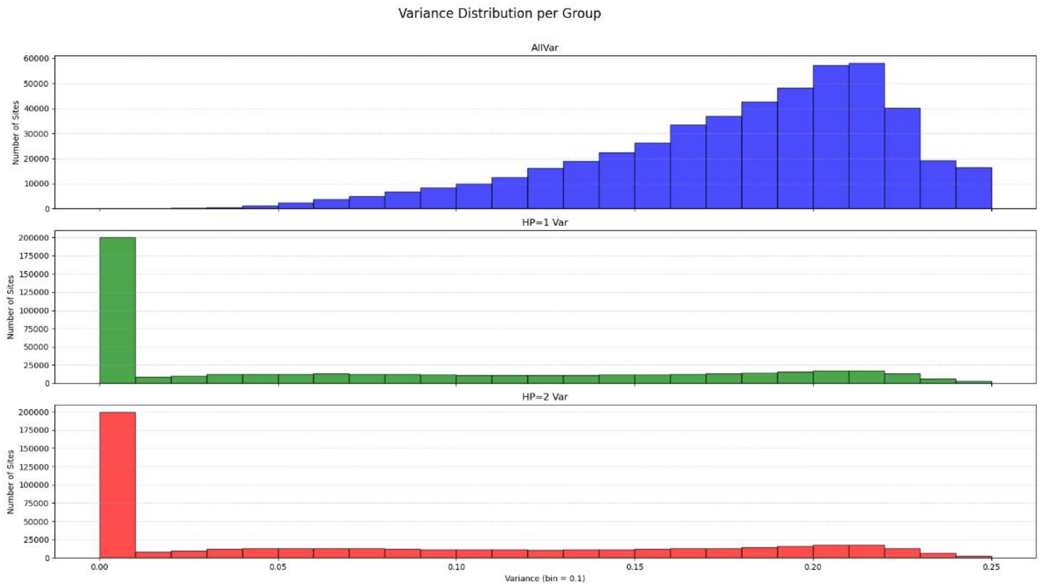


**Figure S7** Variance Distribution Plot of Methylation Sites (Distance = 0.3, Sup_rate =0.3)

***Sup_count***

Under the fixed parameter Distance=0.3, we systematically quantified the number of filtered methylation sites across Sup_count values ranging from 1 to 87, as illustrated in the **Figure S8**.

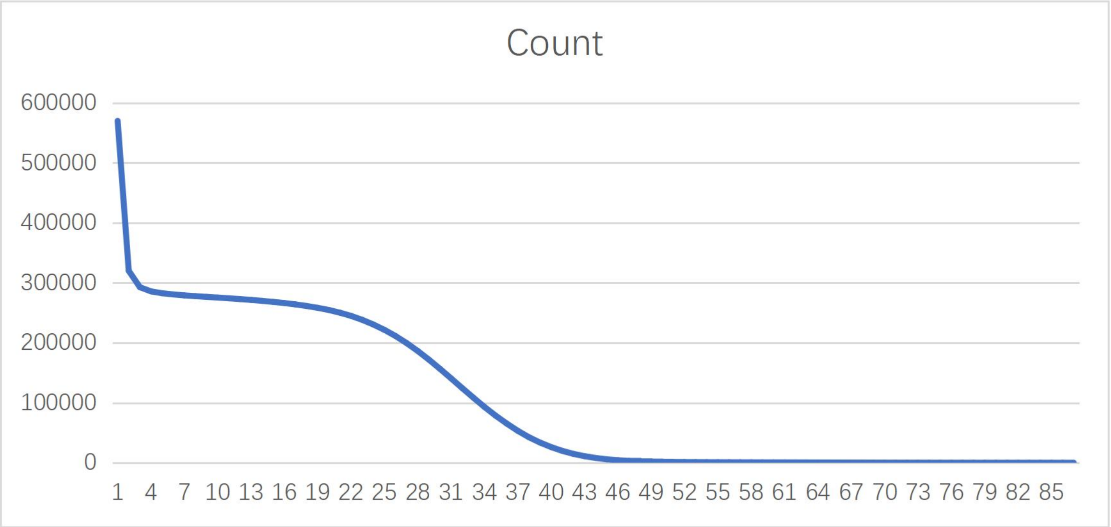


**Figure S8** The number of methylation sites filtered under varying Sup_count thresholds

The figure reveals a distinct inflection point at **Sup_count=4**, where the rate of decline in the number of filtered methylation sites begins to stabilize markedly. This observation justifies setting 4 as the minimum threshold for **Sup_count**.

To determine the upper bound, we adopted the same filtering rationale as in prior analyses by calculating the mean variance of methylation sites retained under varying **Sup_count** values. Key results are summarized in the table below:

From Table S46, it can be observed that starting from **Sup_count=8**, the rate of increase in the mean variance begins to slow significantly. This suggests that the methylation sites filtered by **Sup_count=8** have essentially reached a plateau in terms of haplotype-specific discriminative power.

**Table S45** Mean Variance of Methylation Sites Filtered by Different Sup_count Values.

| Sup_count | variance |
|---|---|
| 2 | 0.186 |
| 3 | 0.1917 |
| 4 | 0.1933 |
| 5 | 0.1939 |
| 6 | 0.1941 |
| 7 | 0.1943 |
| 8 | 0.19442 |
| 9 | 0.19446 |
| 10 | 0.19451 |

Under the optimized parameters (**Distance=0.3**, **Sup_rate=0.3**, **Sup_count=4**), we generated comparative plots using the same methodology as previously described. The results demonstrate enhanced filtering efficacy, with preferential retention of methylation sites exhibiting higher variance values (**Figure S8**).

**Figure S8** displays the variance distribution profiles of methylation sites filtered under different parameter thresholds. Notably, increasing parameter values resulted in a rightward shift of the variance distribution, indicating effective elimination of sites with lower variances. This selective retention pattern suggests that the filtered methylation sites exhibit stronger haplotype-specific characteristics.

To validate the efficacy of these parameters on low-coverage data, we conducted parallel analyses using R9 30× datasets(**Figure S9, Figure S10, Figure S11**).

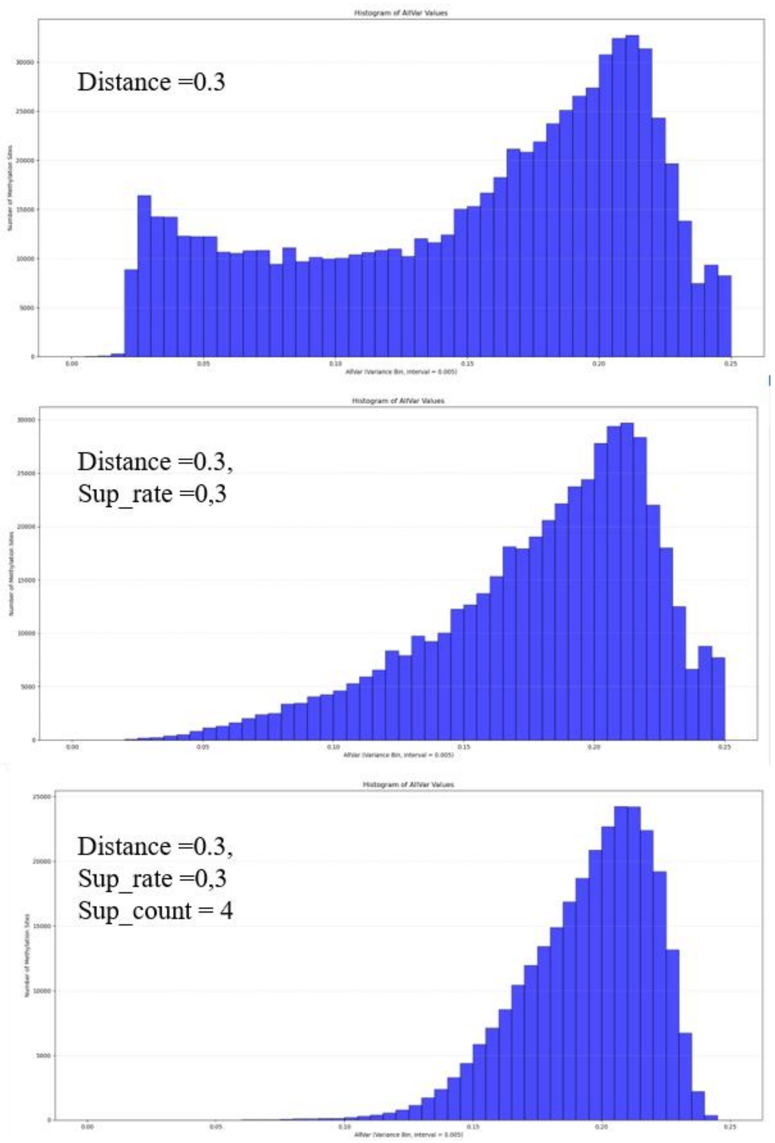


**Figure S8** Variance Distribution Plot of Methylation Sites (R9 80×)

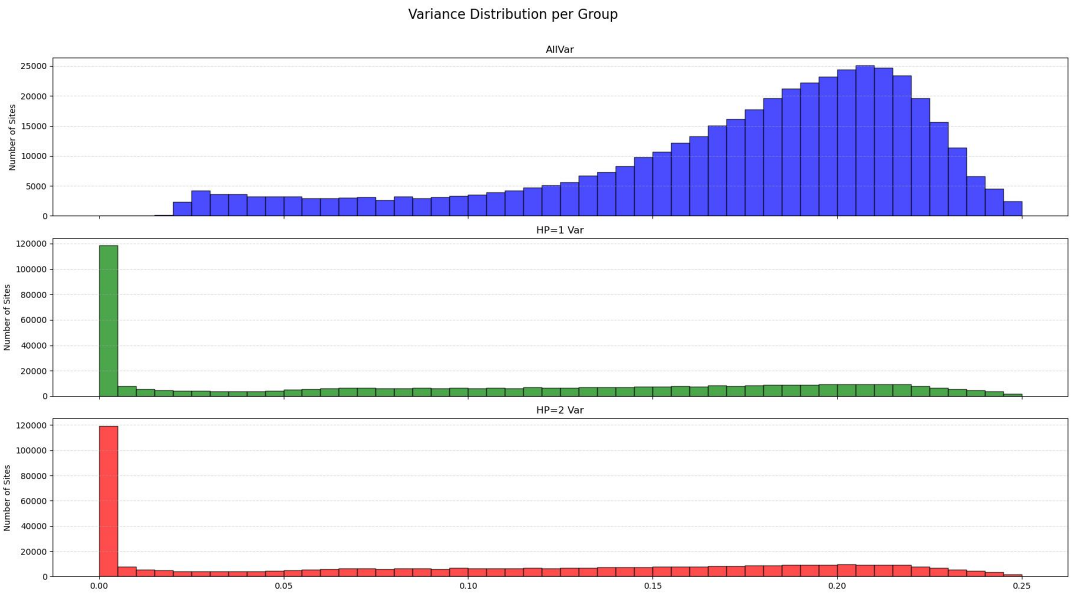


**Figure S9** Variance Distribution Plot of Methylation Sites (R9 30× Distance = 0.3)

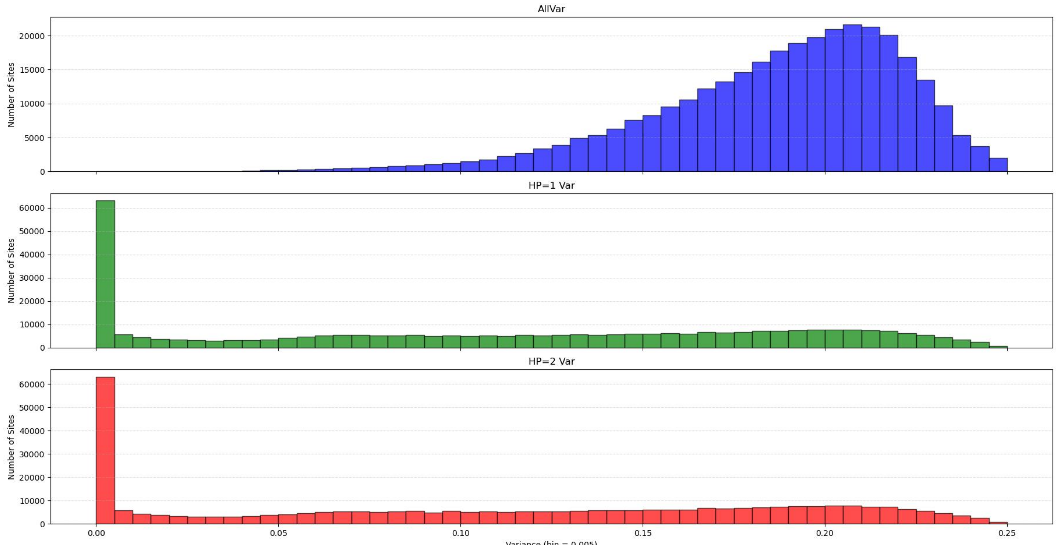


**Figure S10** Variance Distribution Plot of Methylation Sites (R9 30× Distance = 0.3, Sup_rate = 0.3)

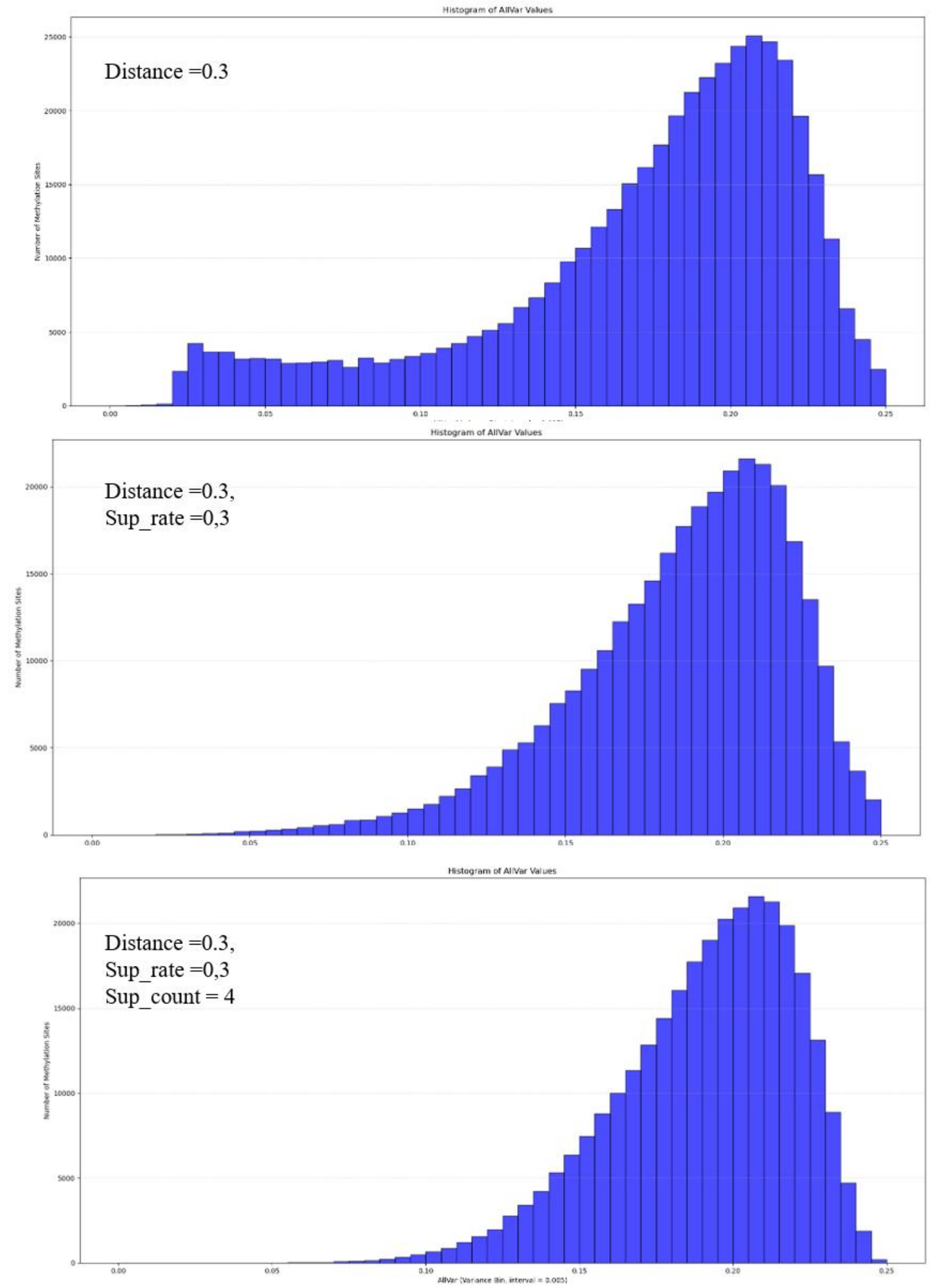


**Figure S11** Variance Distribution Plot of Methylation Sites（R9 30×）

### 5. The Pseudocode

**Algorithm 1** Calculate Consistency Score

```
Input: read_list, where each read contains information on two CpG sites
Output: Methylation CpG locations with higher consistency
function Get_highconsistency_meth(read_list):
    // Iterate through each read and collect CpG site information
    for each read in read_list:
        N_0 = count of CpG sites with (0, 0)
        N_1 = count of CpG sites with (0, 1)
        N_2 = count of CpG sites with (1, 0)
        N_3 = count of CpG sites with (1, 1)
    end for
    // Calculate total number of reads
    total_reads = N_0 + N_1 + N_2 + N_3
    // Calculate the proportion of (0, 0) and (1, 1)
    r = (N_0 + N_3) / total_reads
    // Check for consistency between the two methylation positions
    if |r − (1 − r)| ≥ 0.6 and (N_0 + N_1 + N_2 + N_3) ≥ 10:
        T ← T + 1
    else:
        F ← F + 1
    // Calculate the consistency information for each methylation positions
    X_A = [num(T), num(F)]
    X_B = [num(T), num(F)]
    . . .
    X_N = [num(T), num(F)]
    // Calculate rate for each site
    rate_{X_i} = num(T) / (num(F) + 1)
    // Return methylation positions with high rate values
    return i
```
